# The loss of the pyoverdine secondary receptor in *Pseudomonas aeruginosa* results in a fitter strain suitable for population invasion

**DOI:** 10.1101/2020.05.08.085159

**Authors:** Jaime González, Manuel Salvador, Özhan Özkaya, Matt Spick, Catia Costa, Melanie J. Bailey, Claudio Avignone-Rossa, Rolf Kümmerli, José I. Jiménez

## Abstract

The rapid emergence of antibiotic resistant bacterial pathogens constitutes a critical problem in healthcare and requires the development of novel treatments. Potential strategies include the exploitation of microbial social interactions based on public goods, which are produced at a fitness cost by cooperative microorganisms, but can be exploited by cheaters that do not produce these goods. Cheater invasion has been proposed as a ‘Trojan horse’ approach to infiltrate pathogen populations with strains deploying built-in weaknesses (e.g. sensitiveness to antibiotics). However, previous attempts have been often unsuccessful because population invasion by cheaters was prevented by various mechanisms including the presence of spatial structure (e.g. growth in biofilms), which limits the diffusion and exploitation of public goods. Here we followed an alternative approach and examined whether the manipulation of public good uptake and not its production could result in potential ‘Trojan horses’ suitable for population invasion. We focused on the siderophore pyoverdine produced by the human pathogen *Pseudomonas aeruginosa* MPAO1 and manipulated its uptake by deleting and/or overexpressing the pyoverdine primary (FpvA) and secondary (FpvB) receptors. We found that receptor synthesis feeds back on pyoverdine production and uptake rates, which led to strains with altered pyoverdine-associated costs and benefits. Moreover, we found that the receptor FpvB was advantageous under iron-limited conditions but revealed hidden costs in the presence of an antibiotic stressor (gentamicin). As a consequence, FpvB mutants became the fittest strain under gentamicin exposure, displacing the wildtype in liquid cultures, and in biofilms and during infections of the wax moth larvae *Galleria mellonella*, which both represent structured environments. Our findings reveal that an evolutionary trade-off associated with the costs and benefits of a versatile pyoverdine uptake strategy can be harnessed for devising a Trojan horse candidate for medical interventions.

## Introduction

Microorganisms establish communities where social interactions based on cooperation and competition take place (1). Cooperative strategies, like the synthesis and secretion of essential public goods, usually come with a fitness cost for the cooperative individuals, while carrying a benefit for the whole community (2). These strategies are open to exploitation by cheats, members of the community who do not pay the cost of producing the public good while taking advantage of it, potentially causing the collapse of the population (1). In this context, the use of cheats has been proposed to invade and replace populations of pathogens as a way of treating infections (3). Tailor-made cheats could therefore be used as ‘Trojan horses’ that could keep pathogens resistant to antibiotics at bay by replacing them with sensitive strains (3). The use of cheaters for invasion is, however, hindered because virulence rarely depends on a single essential determinant (4) and by mechanisms preserving cooperation. These include the formation of biofilms, where the diffusion of public goods is limited and restricted to clonal members, thus limiting the emergence of noncooperative individuals (5,6).

Microbial social interactions involving public goods have been widely investigated in the opportunistic human pathogen *Pseudomonas aeruginosa* using the well-studied iron chelator pyoverdine as a model public good (7). Pyoverdine synthesis involves the expression of a large number of genes (8,9) and it generates a fitness cost to producing cells that can be exploited by non-producers (10). Previous attempts at producing a Trojan horse in this system have focused on generating mutants in the biosynthetic pathway of pyoverdine. These non-producers can invade a wildtype population in homogeneous environments (e.g. liquid cultures) but fail to do so in, for instance, animal models (4,11).

In order to overcome that limitation, our study proposes a novel strategy for the generation of Trojan horses. Instead of directly interfering with the synthesis of pyoverdine, we selectively modified its uptake manipulating the pyoverdine receptors. Once pyoverdine binds to iron, the resulting ferripyoverdine is captured by cells mainly through the action of the pyoverdine primary receptor FpvA (12) although a seemingly redundant secondary receptor FpvB can perform the same role (13). In iron-limited conditions, both synthesis and reception through FpvA are pleiotropically coordinated at the transcriptional level through the action of an intricate regulatory circuit involving the activators PvdS and FpvI (14,15) (Fig. 1A). In fact, the binding of ferripyoverdine to the receptor FpvA results in the increased expression of biosynthetic operons (via PvdS) and the receptor gene (via FpvI) in a positive feedback-loop (16–18). This process is only interrupted by the action of the global repressor Fur when the intracellular iron levels are sufficiently high (19).

**Fig. 1.**
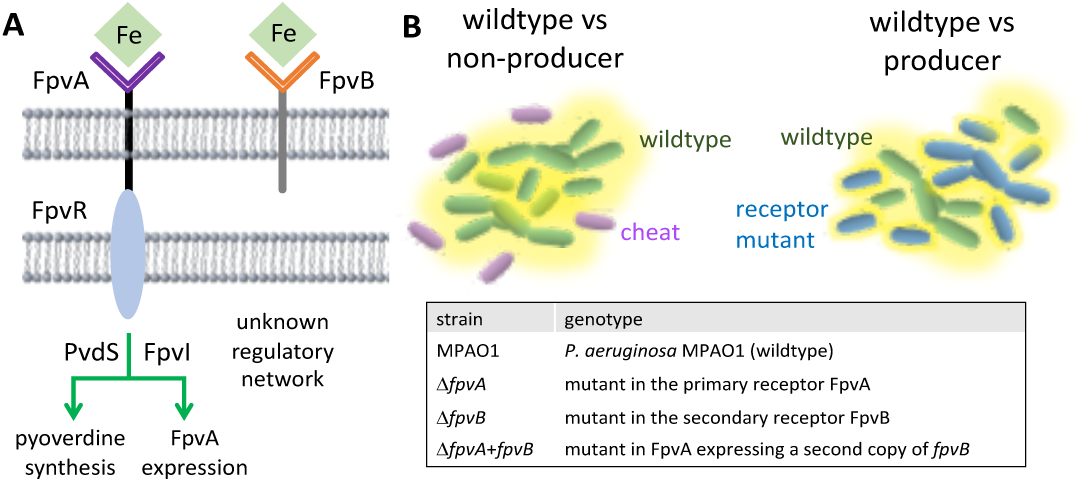
**(A) The regulatory network of the pyoverdine receptors in *P. aeruginosa*.** Ferripyoverdine (green diamonds) bound to the primary receptor FpvA triggers a regulatory signalling cascade resulting in the activation of pyoverdine biosynthesis and FpvA receptor expression. Specifically, pyoverdine-signalling inactivates FpvR (an anti-sigma factor), thereby activating PvdS and FpvI, the sigma factors that control pyoverdine and receptor synthesis, respectively. FpvB participates in the uptake of pyoverdine and its regulatory network is unlinked to the FpvA signalling pathway. Figure inspired by (8). **(B) Outline of the strategy in this study.** Population invasion should be possible with pyoverdine producing strains endowed with optimised cost-to-benefit ratios. The table shows the nomenclature of strains used in this work.

The secondary receptor FpvB, however, is not known to be under the control of the same regulatory network (20) and, therefore, ferripyoverdine binding to it should not lead to an increased pyoverdine synthesis. We hypothesised that altering the expression levels of the primary and secondary receptor could not only affect pyoverdine uptake but also its synthesis due to the interference with the regulatory network. Moreover, it is conceivable that the expression of a secondary, seemingly redundant, receptor could be associated with evolutionary trade-offs and only be beneficial under certain environmental conditions but be costly under others. Thus, interference with pyoverdine uptake could result in strains with a modified ratio of costs (pyoverdine synthesis) and benefits (pyoverdine reception). The specific relationship between those two properties could result in conditional fitness advantages, for example, if the mutants exhibit decreased pyoverdine production and increased cost-effective uptake (Fig. 1B). Such a strain would be able to grow independently (they are public good producers), and have the potential to displace wildtype populations in a variety of scenarios including structured environments.

## Results

### Altered expression of pyoverdine receptors results in different growth and pyoverdine production phenotypes

We monitored growth and pyoverdine production dynamics of *P. aeruginosa* MPAO1 (MPAO1) and the mutants in the primary (Δ*fpvA*) and secondary (Δ*fpvB*) receptors, as well as a mutant that does not produce pyoverdine (Δ*pvdA*) when cultured under iron limitation in the absence or presence of exogenous pyoverdine (Fig. 2 and Fig. S1). In this scenario we define the benefit of a strain by its maximum growth rate achieved which is obtained from the first derivative of the growth curve (see methods). Maximum growth rate is a proxy of the physiological/metabolic efficiency of a cell when protein production is at the steady state. Similarly, the cost of a strain is defined as the pyoverdine produced up to the time in which the maximum growth rate is achieved. This reflects a metabolic investment that greatly conditions the growth rate at the early stages of growth of the strains tested.

**Fig. 2.**
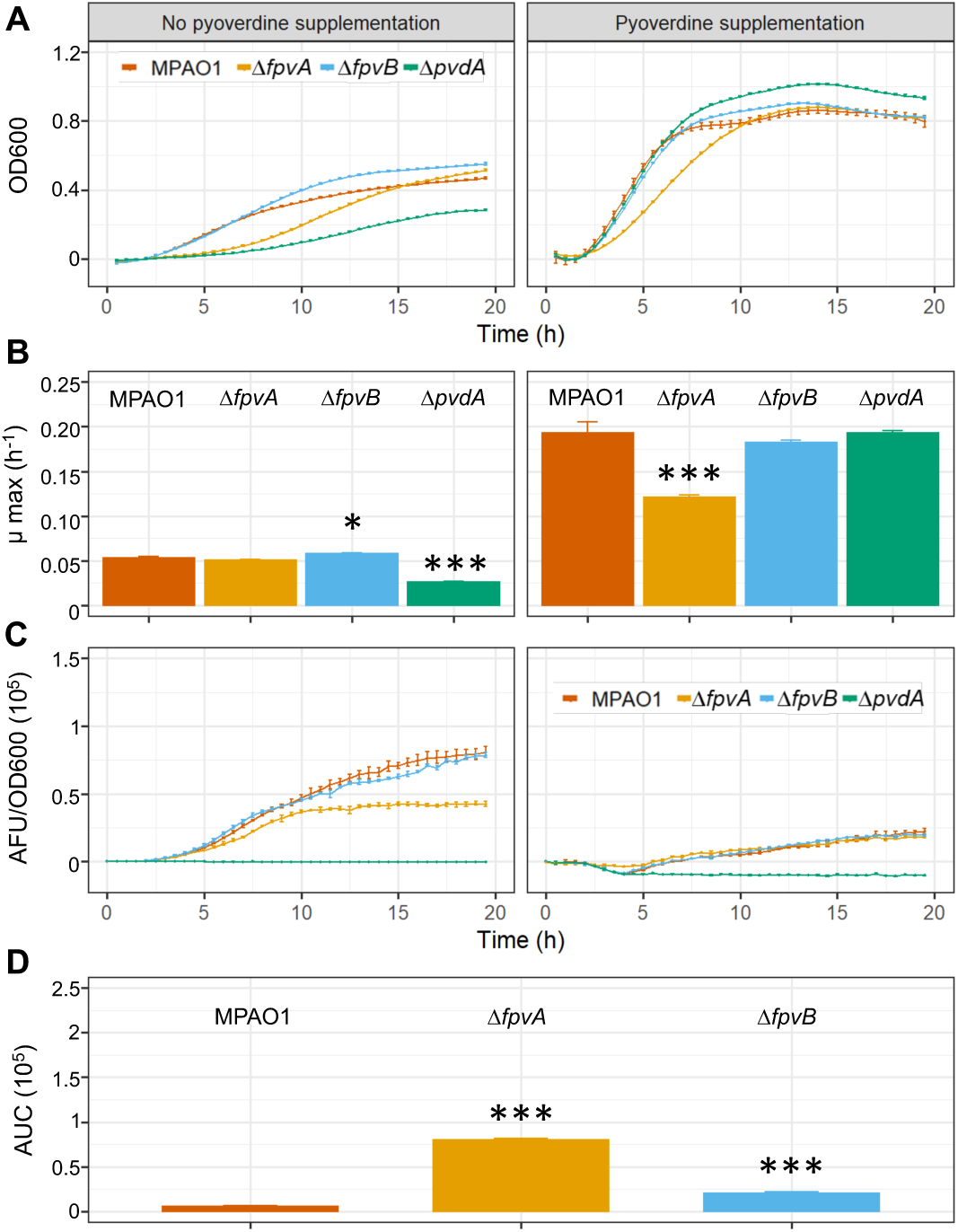
Kinetics of bacterial growth and pyoverdine production. (A) Growth kinetics of MPAO1 and mutants in iron-limited CAA in the absence (left column) and presence (right column) of exogenous pyoverdine. (B) Maximum growth rates of the strains obtained from the analysis of growth kinetics. (C) Pyoverdine production per cell over time determined as the fluorescent signal of the culture divided by the optical density at 600 nm (OD600). (D) Pyoverdine accumulated in cultures of each of the strains until reaching the maximum growth rate in the absence of exogenous pyoverdine. Error bars represent standard error of the mean from three biological replicates. Asterisks show significant differences compared to MPAO1 in a One-way ANOVA test complemented with a Dunnet post-hoc test. Growth curves and their first derivatives are shown in Fig S1.

All strains exhibited similar maximum growth rates (ranging from 0.0505 ± 0.0013 to 0.0578 ± 0.0126 h^-1^) with the exception of the pyoverdine deficient Δ*pvdA* mutant, which had a significantly reduced growth rate (0.026 ± 0.001 h^-1^). Moreover, the mutant Δ*fpvA* lacking the primary receptor showed a longer lag phase compared to the other producers (Figs. 2A and B left column; Fig. S1). When supplemented with exogenous pyoverdine, which should alleviate the burden of production, all the strains exhibited similar growth rates (ranging between 0.181 ± 0.002 and 0.192 ± 0.012 h^-1^) and kinetics except for Δ*fpvA*, which had a significantly reduced growth rate (0.121 ± 0.00176 h^-1^) and an extended lag-phase (Fig. 2A and B right column; Fig. S1). Altogether, these results suggest that FpvA is essential for optimal pyoverdine uptake rates, and that the secondary receptor FpvB can only partly compensate for the lack of FpvA, leading to a suboptimal pyoverdine uptake.

Next, we inspected pyoverdine production in all strains in both conditions. Under iron-limited conditions, the Δ*fpvA* mutant exhibited a lower pyoverdine synthesis per cell but a larger investment prior to reaching the maximum growth rate, suggesting that the primary receptor plays an important role in both efficient uptake and coordination of pyoverdine synthesis (Fig. 2C and D; Fig. S1). Interestingly, the wildtype strain performed best when considering the amount of pyoverdine produced until reaching the maximal growth rate (Fig. 2D), which indicates that having two different pyoverdine receptor types (FpvA and FpvB) lead to the most economical cost-to-benefit ratio.

### Pyoverdine production is affected by the presence of gentamicin

Previous studies have reported that the costs and benefits linked to pyoverdine biology in *P. aeruginosa* are condition-dependent (21). In particular, the cost of producing pyoverdine increased when cells were exposed to environmental stressors such as sublethal concentrations of gentamicin (22).

Since aminoglycosides are commonly used to treat *P. aeruginosa* infections (23) we inserted the *aacC1* resistance gene at the attTn7 site of all the strains and investigated synthesis of pyoverdine and reception dynamics in the presence of gentamicin (Fig. 3 and Fig. S2). In iron-limited conditions gentamicin caused longer lag phases (Fig. 3A) and a general decrease in growth rates (Fig. 3B), showing that gentamicin acts as a stressor despite strains being resistant to it. Crucially, gentamicin altered the rank of growth rates with Δ*fpvB* (0.038 ± 0.0016 h^-1^) and Δ*fpvA* (0.032 ± 0.001 h^-1^) showing higher growth capacities than MPAO1 (0.022 ± 0.001 h^-1^). Addition of pyoverdine led to an improved growth and lag phase reduction in all the strains (Fig. 3A and B). But also here Δ*fpvB* growth rate was higher (0.145 ± 0.003 h^-1^) compared to Δ*fpvA* (0.109 ± 0.0022 h^-1^) and MPAO1 (0.12 ± 0.004 h^-1^) (Fig. 3B).

**Fig. 3.**
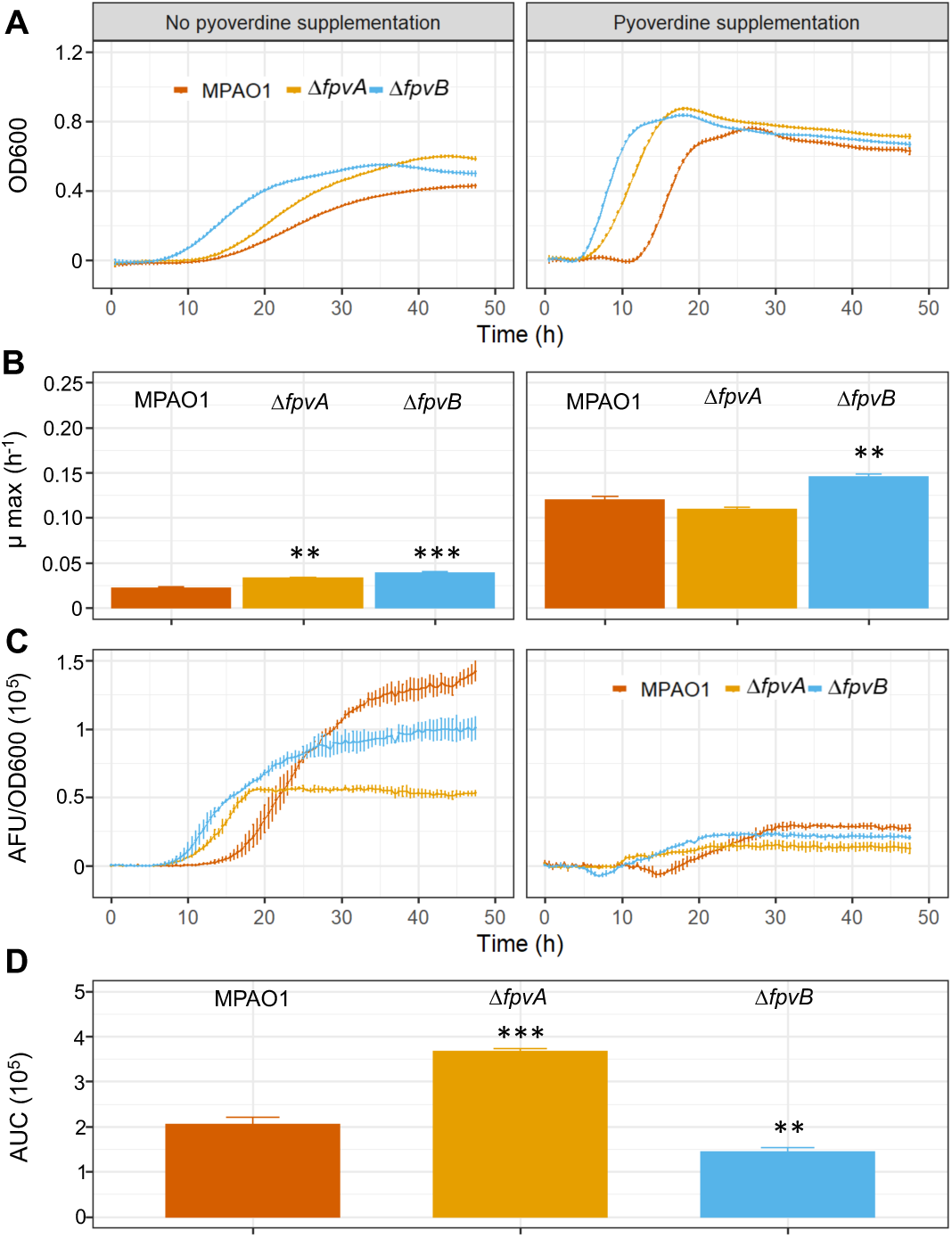
Kinetics of bacterial growth and pyoverdine production in the presence of gentamicin. (A) Growth kinetics of MPAO1 and mutants in iron-limited CAA supplemented with gentamicin in the absence (left column) and presence (right column) of exogenous pyoverdine. (B) Maximum growth rates of the strains obtained from the analysis of growth kinetics. (C) Kinetics of pyoverdine production per cell. (D) Pyoverdine accumulated in cultures of each of the strains until reaching the maximum growth rate. Error bars represent standard error of the mean from three biological replicates. Asterisks show significant differences compared to MPAO1 in a One-way ANOVA test complemented by a Dunnett post-hoc test. Growth curves and their first derivatives are shown in Fig S2.

Pyoverdine production per cell over time varied fundamentally between cells in iron-limited but not in pyoverdine supplemented medium (Fig. 3C). Under iron limitation, Δ*fpvB* started producing pyoverdine first, while MPAO1 showed a substantially delayed production, which is reduced in Δ*fpvA*. The advantage of Δ*fpvB* over the other strains is mirrored in the production levels up to maximum growth rate (Fig. 3D), where Δ*fpvB* performed most economically followed by MPAO1 and Δ*fpvA*. When comparing across conditions (with and without gentamicin), we found that MPAO1 produced 30 ± 2.7 times more pyoverdine (as AUC ratio values) to reach its maximum growth rate with gentamicin, while the ratio was only slightly higher for Δ*fpvA* (2.4 ± 0.004) and Δ*fpvB* (2.58 ± 0.17), respectively (Fig. S3). Taken together, these results show that Δ*fpvB* performs best under antibiotic stress, suggesting that having a secondary pyoverdine receptor involves a costly evolutionary trade-off. Thus, the Δ*fpvB* mutant could be a promising candidate for a Trojan horse.

### *ΔfpvB* expresses the primary receptor FpvA earlier than MPAO1 in the presence of gentamicin

We monitored the expression dynamics of the two pyoverdine receptors using transcriptional fusions in which the corresponding promoters were cloned in a Tn7 transposon (Gm^R^) for chromosomal delivery right upstream of a promoterless mCherry gene. Both constructions were independently integrated in single copy in the attTn7 site of the strains MPAO1, Δ*fpvA* and Δ*fpvB*. This allows for the monitoring of gene expression levels even in mutants unable to express the genes under study.

The expression kinetics of the primary receptor FpvA differs between MPAO1 and Δ*fpvB*. In the absence of gentamicin, MPAO1 triggered the expression of the primary receptor FpvA earlier and also displayed higher transcriptional levels compared to the Δ*fpvB* strain (Fig. 4A left panel). When gentamicin was added to the cultures, the expression signal for the FpvA receptor increased overall in both the MPAO1 and Δ*fpvB* strains, but Δ*fpvB* transcribed *fpvA* much earlier than MPAO1 (Fig. 4A right panel). Transcriptional levels of the secondary receptor *fpvB* were low for all strains except in the mutant Δ*fpvA* that seems to compensate for the lack in the primary receptor by increasing the expression of the secondary (Fig. 4B). These results highlight that Δ*fpvB* has a markedly different expression pattern of the primary receptor FpvA than MPAO1, characterized by a reduced expression level combined with an earlier onset of expression in the presence of the environmental stressor. This altered expression pattern likely contributes to the fitness advantage of the Δ*fpvB* mutant under gentamicin exposure.

**Fig. 4.**
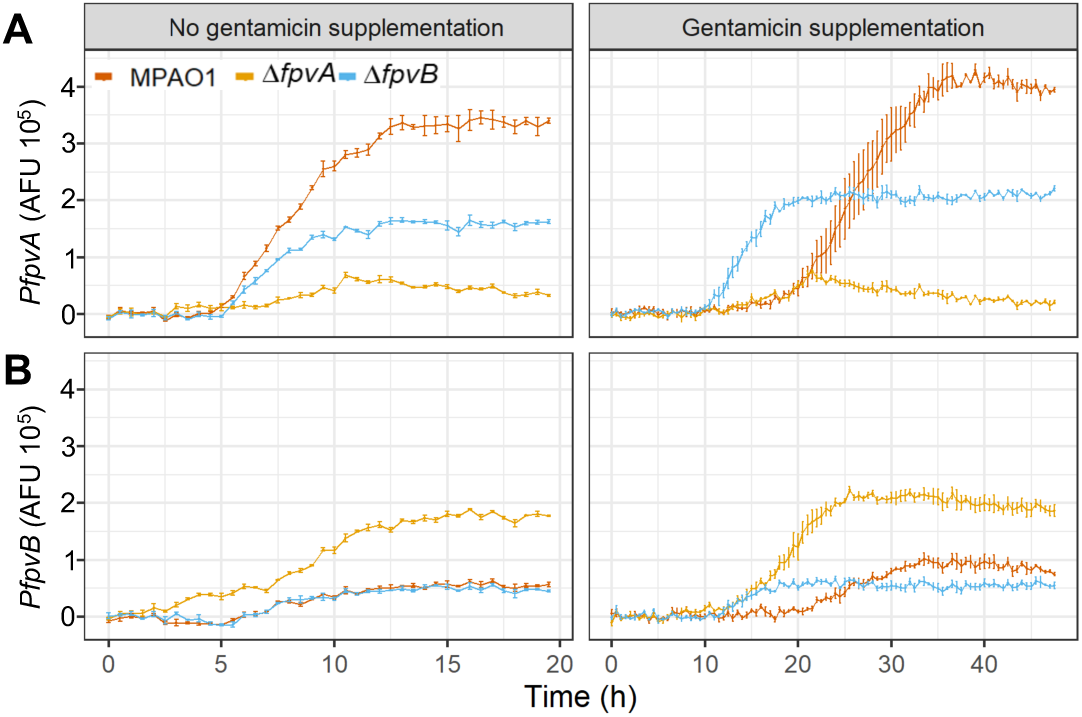
Analysis of the transcriptional activity of the promoters of *fpvA* and *fpvB* in iron limited CAA. Transcriptional fusions between the *fpvA* (A) or *fpvB* (B) promoters and mCherry were delivered into the attTn7 of the strains indicated in the bars and tested in the absence (left columns) and presence (right columns) of gentamicin. The curves represent the fluorescent signal per cell over time. Error bars represent standard error of the mean from three biological replicates.

### The conditional expression of FpvB tunes growth and pyoverdine production

Our previous results show that the loss of FpvB renders a strain with a higher growth rate and lower pyoverdine investment in the presence of gentamicin, suggesting that the competition between the two receptors controls the trade-offs between the costs and benefits associated to the public good. We therefore tested the role of the secondary receptor FpvB as a dial to tune growth dynamics in a strain that possesses both receptors (MPAO1) and a strain that only has FpvB (Δ*fpvA*) (Fig. 5; Figs. S4 and S5). To this end, a recombinant genetic construct containing *fpvB* under the regulation of a strong constitutive promoter (14F; (24)) was introduced as an extra copy in the genome, including the gentamicin resistance cassette *aacC1*.

**Fig. 5.**
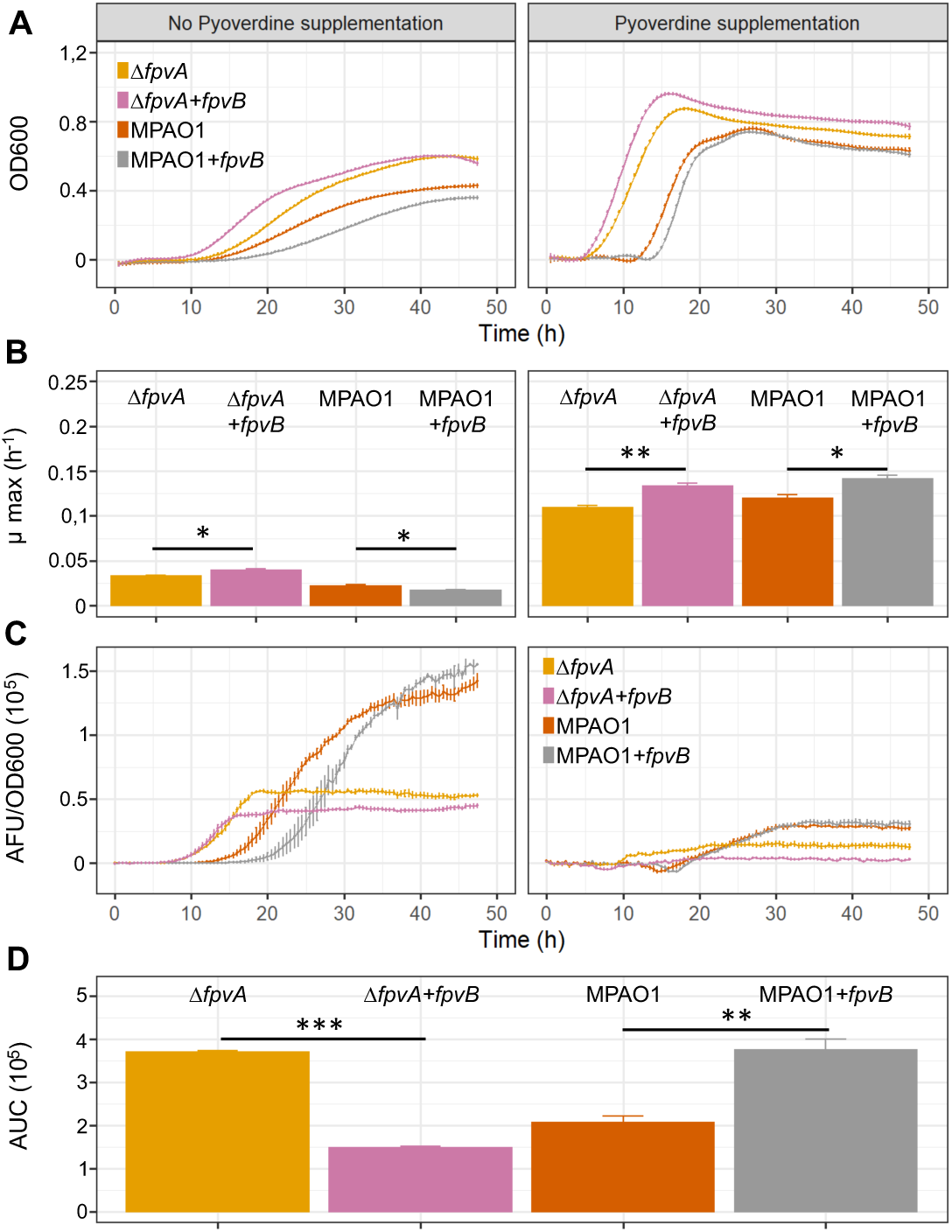
Kinetics of bacterial growth and pyoverdine production in strains overexpressing FpvB in the presence of gentamicin. (A) Growth kinetics of MPAO1 and Δ*fpvA* expressing or not an extra copy of *fpvB* in iron-limited CAA supplemented with gentamicin in the absence (left) and presence (right) of exogenous pyoverdine. (B) Maximum growth rates of the strains obtained from the analysis of growth kinetics in the same conditions. (C) Kinetics of pyoverdine production per cell. (D) Pyoverdine accumulated in cultures of each of the strains until reaching the maximum growth rate. Error bars represent standard error of the mean from three biological replicates. Asterisks show significant differences compared the corresponding ‘empty’ control in a t-test. A complete set of growth curves, their first derivatives and pyoverdine production kinetics is shown in Fig S5).

In the presence of gentamicin, the overexpression of FpvB was advantageous for Δ*fpvA* which reached a higher maximum growth rate both in iron-limited conditions (0.039 ± 0.001 vs 0.032± 0.0010 h^-1^) and when exogenous pyoverdine was added (0.13 ± 0.002 vs 0.109 ± 0.002 h^-1^) (Fig. 5A and B; Fig. S5). This shows that FpvB can alleviate iron shortage associated with the lack of the primary receptor FpvA. In stark contrast, overexpression of FpvB in MPAO1 had the opposite effect and induced a longer lag phase (Fig. 5A) and decreased the maximum growth rate (0.017 ± 0.0006 vs 0.022 ± 0.0009 h^-1^) (Fig. 5B). This growth deficiency was partly mitigated with the addition of exogenous pyoverdine (Fig. 5A and B; Fig. S5). These findings support the view that FpvB expression is associated with substantial fitness costs in the presence of a functional FpvA receptor under gentamicin stress. The fitness trade-off associated with FpvB expression is also reflected in the pyoverdine production profiles (Fig. 5C and D), where FpvB overexpression leads to a reduction of pyoverdine per cell required to reach maximum growth rate in the Δ*fpvA* strain, but leads to the opposite pattern in MPAO1 (Fig. 5D).

### The *ΔfpvB* mutant rapidly invades MPAO1 populations from rare under gentamicin treatment

Next, we examined whether the growth properties uncovered are predictive of population dynamics when strains with different pyoverdine uptake strategies compete directly in batch liquid cultures. To this end, gentamicin resistant GFP and RFP tagged strains were mixed in a 0.85:0.15 (MPAO1: mutant) initial proportion and transferred to iron limited CAA, with and without gentamicin supplementation. Population dynamics were monitored using flow cytometry (Fig. 6 for frequencies; see Fig. S6 for relative fitness) and we confirmed that gentamicin remained at high levels in all of the competitions (Fig. S7).

**Fig. 6.**
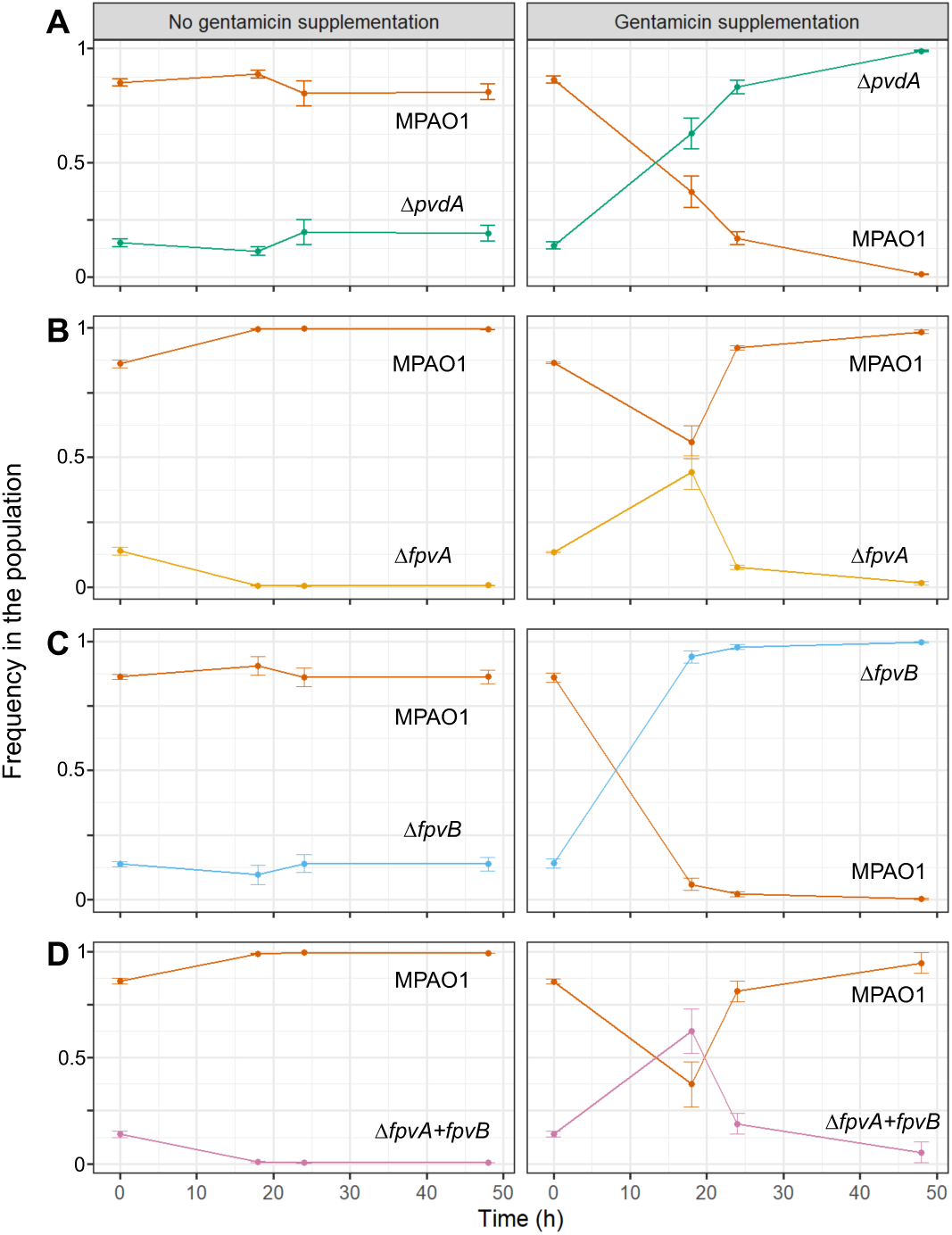
Dynamics of bacterial competition in batch cultures in iron limited CAA in the presence or absence of gentamicin. Each plot represents the frequencies of the two competing strains over time in the absence (left column) or presence (right column) of gentamicin. Initial proportions were 0.85:0.15 of, respectively, MPAO1 and the mutants Δ*pvdA* (A), Δ*fpvA* (B), Δ*fpvB* (C) and Δ*fpvA+fpvB* (D). Frequencies were determined analysing 5·10^4^ event counts in a flow cytometer. Error bars represent standard error of the mean from three biological replicates with alternate fluorescent tags (GFP and RFP; results of one competition were confirmed determining cfus in a plating assay and are shown in Fig. S8).

In the absence of gentamicin, we found that none of the mutants could invade MPAO1 populations (Fig. 6). While Δ*fpvA* mutants went completely extinct, the Δ*fpvB* and Δ*pvdA* mutants could co-exist with MPAO1. The latter likely due to its capacity to cheat on pyoverdine produced by MPAO1. In the presence of gentamicin, the strains Δ*fpvA* and Δ*fpvA*+*fpvB* still went extinct although showed an increase in frequency in the first 18 h of the experiment (Fig. 6). This is likely the result of MPAO1 being the fittest strain when enough pyoverdine was accumulated, which allowed to overcome the initial disadvantage. Conversely, the Δ*fpvB* and Δ*pvdA* mutants experienced huge fitness advantages under antibiotic stress and drove MPAO1 to the verge of extinction over the 48 hours competition. The spread of the cheating Δ*pvdA* mutant matches the results from a previous study (22), showing that pyoverdine is especially costly under stressful conditions, which makes cheating particularly profitable. The rapid and consistent spread of the Δ*fpvB* mutant supports our hypothesis that expression of this secondary receptors is particularly costly under antibiotic stress and its deletion allows mutants to invade from rare.

### The ΔfpvB mutant is a suitable Trojan horse in structured biofilms and in an animal model

The presence of spatial structure limits the diffusion of public goods such as pyoverdine, thereby leading to changes in pyoverdine distribution and uptake rates. This in turn can feed back on the dynamics between competing strains in microbial populations (25,26). To test whether *ΔfpvB* mutants can also invade MPAO1 populations in spatially structured environments, we first repeated the competition experiments by inoculating mixtures of MPAO1 and the different mutants (0.85:0.15 MPAO1:mutant initial mixing ratio) into a multichannel chamber (see Methods) in iron limited CAA, where cells attach to the surface and form structured biofilms. After 48 hours, the resulting biofilms were imaged with a confocal microscope and quantitative information was obtained by integrating the area in the image corresponding to each colour. As in liquid cultures, we found that both Δ*pvdA* and Δ*fpvB* dominated the biofilm in the presence of gentamicin, driving MPAO1 to almost complete extinction (Fig. 7A). In contrast, Δ*pvdA* (in the absence of gentamicin) and Δ*fpvA* mutants were able to co-exist with MPAO1 but could not dominate it. Unlike in liquid cultures, these latter set of mutants could increase in frequency in the biofilms (Fig. 7A).

**Fig. 7.**
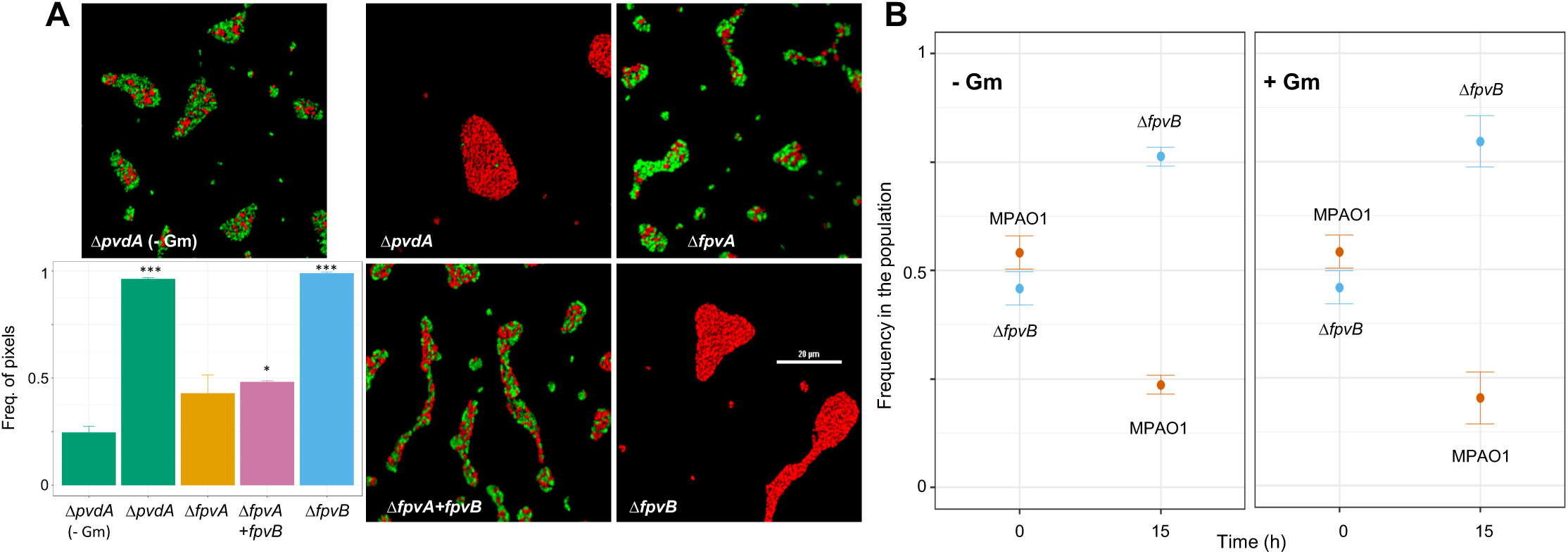
Dynamics of bacterial competition in structured environments. (A) *In-vitro* biofilm formation in competition assays between MPAO1 and the mutants indicated. All pictures and bars correspond to cultures with gentamicin except the labelled as -Gm. Bars represent pixel frequency of red pixels (mutant strains). Each image is representative of a set of 10 randomly selected areas (selected using phase contrast to avoid bias) for each condition, taken for each of the three biological replicates, from which standard error bars were generated. Scale bars equal 20 μm (see Figs. S9 and S10 for larger pictures). Asterisks show significant differences compared to the competition MPAO1-Δ*pvdA* (-Gm) in a One-way ANOVA test complemented by a Dunnett post-hoc test. (B) Competition assays in *G. mellonella.* A population of 10^2^ to 10^3^ cells containing an initial proportion of 0.5:0.5 MPAO1 to Δ*fpvB* was injected in the larvae in the absence (left panel; -Gm) and presence (right panel; +Gm) and let grow for 15 hours. Subsequently, animals were culled and strain frequencies quantified. Error bars represent standard error of the mean for 3 biological replicates with alternate fluorescent tags (GFP and RFP).

We then used *Galleria mellonella* larvae as a model to reproduce the spatial structure *P. aeruginosa* would face in a host during an infection (27). We infected larvae with mono- and mixed cultures in the presence or absence of gentamicin following a standard protocol of infection (28), using 10^2^-10^3^ cells as the inoculum. Single strain infections were performed with MPAO1, Δ*fpvA* and Δ*fpvB* in order to obtain survival curves of the animal host in the presence and absence of gentamicin. Independently of the presence of the antibiotic, all larvae died within 22 hours when infected with bacteria, and there were no significant differences between the two treatments (Fig. S11). We then tested whether Δ*fpvB* can also dominate MPAO1 in this host environment by establishing mixed infection with the two competing strains. We found that this was indeed the case (Fig. 7B). Starting from an initial proportion of 0.46 ± 0.04, Δ*fpvB* increased to frequencies of 0.76 ± 0.02 (without gentamicin) and 0.80 ± 0.06 (with gentamicin) in only 15 hours. These results show that even in the animal model the Δ*fpvB* mutant can invade wild type populations.

## Discussion

Selective interventions steering the evolutionary dynamics of microbial communities with the main goal of reducing antibiotic usage are at the core of microbial evolutionary medicine (29). In this work we have investigated the effect of interfering with the reception of pyoverdine in *P. aeruginosa* as a way to engineer strains with the potential of invading wildtype populations. Our results show that versatile pyoverdine uptake strategies are beneficial in the absence of a stressor, possibly reflecting evolutionary adaptations to environments with limited and/or fluctuating iron availabilities, but are costly under antibiotic stress. This fitness trade-off can be used to engineer a highly invasive strain, potentially useful as a Trojan horse. Among the mutants tested, Δ*fpvB* that lacks the secondary receptor outcompeted MPAO1 due to a lower cost of pyoverdine production while modulating high receptor-linked benefits when in the presence of the antibiotic gentamicin used as an environmental stressor. In this condition all strains responded to gentamicin by increasing pyoverdine production, which resulted in a significant delay in growth as well as a lower growth rate in MPAO1. It required 30 times more pyoverdine to reach the maximum growth rate compared to 2.5 times in the other strains including Δ*fpvB*. The ability of gentamicin to increase the expression of some genes involved in pyoverdine and pyochelin (secondary iron chelator) synthesis even when iron is available has been described previously (30,31) but the precise mechanism by which this takes place is unknown.

Our results offer insights into the properties of the secondary receptor: supplementing with pyoverdine support that growth at the expense of FpvB was suboptimal, likely due to a lower efficiency in pyoverdine uptake via FpvB – both receptors share a 54% amino acid sequence similarity (38% identity) – (13,32). However, FpvB can be used as a dial to control growth kinetics and its overexpression mitigated the growth defects of a FpvA mutant but was detrimental for MPAO1, which exhibited a delayed growth and required a higher pyoverdine production to reach its maximum growth rate. The differences can be attributed to the molecular intervention impairing the regulatory network that directly links pyoverdine synthesis and reception, a scenario where Δ*fpvB* had a selective advantage. Actually, pyoverdine receptors generate a cost to the cells (33) and mutations in FpvB have been detected in clinical isolates obtained from cystic fibrosis patients (34). Our results show that FpvB is generally advantageous under iron limitation, but generates a big cost to the cell in stressful conditions. In these conditions, a mutant lacking the secondary receptor is fitter than the wildtype and therefore more resistant, becoming an ideal candidate for outcompeting MPAO1.

The growth profiles in monocultures were predictive of the outcomes of the competition experiments in liquid cultures apart for Δ*pvdA*, which grew badly in isolation, but performed well in co-culture with MPAO1 in the absence of gentamicin. This pattern is compatible with cheating, where pyoverdine non-producers obtain benefits by exploiting producers, especially at low initial frequency (35). When competing against receptor mutants, MPAO1 had equal fitness compared to Δ*fpvB*, but outcompeted Δ*fpvA*, showing that FpvA is essential for efficient iron acquisition in iron-limited medium. However, when gentamicin was added, both Δ*fpvB* and Δ*pvdA* drove MPAO1 to almost extinction, with the latter finding being in agreement with a previous study (22). Thus, both mutants could be used as Trojan horses although we predict that Δ*fpvB* is the more promising candidate because it does it faster and, importantly, because it does not depend on the presence of a pyoverdine producer for iron acquisition. Moreover, our results show that gentamicin is essential for Δ*pvdA* invasion, a finding that could explain the apparent discrepancy in the selective advantage of pyoverdine-deficient mutants reported in previous studies. Specifically, pyoverdine mutants were unable to invade *P. aeruginosa* wild type strains in infections of the animal models *G. mellonella* and *C. elegans* where no antibiotics were administred (4,11), while they could efficiently displace wild type pyoverdine producers in the lungs of patients with cystic fibrosis, who typically undergo intense antibiotic treatments (36).

The strain Δ*fpvB* dominated competitions on a surface in the presence of gentamicin. In fact, in this condition all mutants increased their frequency against MPAO1, which suggests that pyoverdine is not a relevant driver of the population dynamics in this condition. This could be the result of additional factors, such as the anoxic environment found in biofilms, which increase iron availability (37). In addition, Δ*fpvB* also invaded MPAO1 in infections of the animal model *G. mellonella*, in which *P. aeruginosa* shows a virulence that correlates with that observed in mice (38). Unlike in the previous scenarios, the presence of gentamicin in the animal did not lead to observable differences in the growth of the strains and, as a result, their pathogenicity was not affected. This is consistent with a previous meta-analysis of different animal models showing that the lack of pyoverdine modestly reduces virulence *in vivo*, particularly in invertebrates (39). As in biofilms, iron acquisition via pyoverdine might not be essential for colonising the larvae. However, this does not mean that competition-cooperation dynamics are not taking place and, in fact, our results confirm that the pyoverdine system confers a fitness advantage for growth in the animal (11). This suggests that the *G. mellonella* can at least partly replicate the stress conditions that impact the *in vitro* evolutionary dynamics based on pyoverdine (40,41).

Our findings highlight the potential of manipulating pyoverdine receptors to obtain Trojan-horse strains with the ability to dominate a population under specific conditions. Out of the strains characterised, Δ*fpvB* was capable of invading MPAO1 both in structured and unstructured environments including an animal model. It did not require the presence of exogenous pyoverdine for growth, since it could still produce it, therefore overcoming the limitations of strains deficient in pyoverdine production. This mutant could constitute the basis of a bacterial vehicle for the deployment of traits of interest in wildtype populations.

## Materials and Methods

### Strains

All *Pseudomonas* strains were obtained from a transposon mutant library and are described in the supplementary materials. *E. coli* DH5α and One Shot™ PIR2 (Thermofisher) were required for standard genetic manipulations and plasmid maintenance (Tables S1 and S2).

### Culture conditions and real time monitoring of pyoverdine, OD and FP production

Lysogeny broth (LB) contained 10 g peptone, 5 g yeast extract and 10 g NaCl per litre of media. For agar plates, 15 g·L^-1^ of agar were added. Iron-limited casamino acid media (CAA) was prepared using 5 g vitamin free casamino acids, 1.18 g K_2_HPO_4_.3H_2_O, 0.25 g MgSO_4_·7H_2_O, per litre of distilled H_2_O, supplemented with 200 µg·mL^-1^ of human apo-transferrin, and 20 mM sodium bicarbonate. CAA pH was adjusted to 7 and buffered using 50 mM HEPES. Apo-transferrin is used to avoid iron sequestration by secondary chelators generated by *P. aeruginosa*, while sodium bicarbonate is required for its activity (7). Pseudomonas Isolation Agar plates were prepared following the manufacturer (Fisher Scientific, UK) instructions.

Media were supplemented with 20 µg·mL^-1^ of gentamycin, 30 µg·mL^-1^ of chloramphenicol, 50 µg·mL^-1^ of kanamycin and 150 µg·mL^-1^ of ampicillin when required. To supplement bacterial growth with pyoverdine, *P. aeruginosa* Δ*pchE* was grown in CAA for 48 hours and the supernatant was filter-sterilized three times. The use of this mutant avoided production of the secondary iron chelator pyochelin (42). In the required condition, this supernatant was added (10% vol/vol) to fresh CAA. All reagents were obtained from Sigma-Aldrich (UK) unless stated otherwise.

For all monocultures, cells were taken from single clone frozen stocks and grown overnight at 37°C 200 rpm in 5 mL LB supplemented with antibiotics if required. The following day, cells were transferred (1%) to fresh LB in standard 24 well plates and grown to mid-exponential phase (BMG-Clariostar, 37°C, 500 rpm). Once cells reached mid-exponential phase, they were washed 3 times in phosphate saline buffer (PBS), and OD600 normalized to 1 (Thermo Scientific Evolution 60S UV-Visible Spectrophotometer). Cells were then diluted to OD600 0.01 in CAA (supplemented when required with pyoverdine and/or gentamicin) and grown in standard 96 well plates (BMG-Clariostar, 37°C, 500 rpm). OD600, pyoverdine fluorescence (450 nm(Ex)/490 nm(Em)) and mCherry fluorescence (587 nm(Ex)/610 nm(Em)) were measured every 30 minutes.

### General molecular biology techniques

DNA amplification was performed using the Q5 DNA polymerase kit (New England Biolabs, USA) and the corresponding primers (Table S3) following the recommendations of the manufacturer.

Otherwise stated, plasmid construction was carried out using standard digestion and ligation procedures. The general protocol involved plasmid extraction using the QIAprep spin mini-prep kit (Qiagen, UK). 1 µg of plasmid was mixed with 10 U of the corresponding restriction enzymes and the corresponding amount of 1X restriction buffer in final 50 µL volume (New England Biolabs, USA). Reactions were incubated at least for one hour at the recommended temperature. After digestion, fragments were purified using the QIAquick PCR Purification Kit (Quiagen, UK) or, QIAquick Gel Extraction Kit (Qiagen, UK) following the instructions of the manufacturer. The resulting plasmids were chemically transformed into DH5α or One Shot™ PIR2 chemically competent *E. coli* cells (Invitrogen, UK). After transformation, cells were resuspended in 1 mL of LB and grown for one hour (37°C, 200 rpm) before plating on LB agar supplemented with the required antibiotics. Descriptions of the plasmid construction strategies as well as lists of primers, strains and plasmids used in this study can be found in the supplementary methods.

### Tn-7 chromosomal integration

Donor (One Shot™ PIR2 *E. coli* - for R6K plasmid replication-), recipient (*P. aeruginosa* strains) and helper strains (*E. coli* DH5α carrying pRK600 and pTns-1 plasmids) were grown overnight in 5 mL of LB supplemented with the required antibiotics. The helper and donor strains were re-inoculated (1%) in 1 mL of fresh LB and grown until 0.4-0.5 OD at 37°C. The recipient strains were re-inoculated (10%) in 1 mL of fresh LB and grown for the same amount of time than the helper and donor strains at 42°C with no agitation. The cells were then washed twice with 1 mL of LB to remove antibiotics. 100 µL of each strain were mixed, centrifuged, re-suspended in 30 µL of LB and plated onto an antibiotic-free LB agar plate. After 6 hours at 37°C, the cells were recovered from the plate in 1 mL of LB. The cell pellet was harvested, resuspended in 30 µL of PBS and plated on Pseudomonas Isolation Agar with gentamycin to select for bacteria carrying the genetic construction. Tn7 presence was confirmed by colony PCR using Tn7RFW and glmsDownREV primers (Table S3) or whole genome sequencing in the case of FpvB insertion.

### *In vitro* competition studies

Once grown and OD600 normalized to 1, cells were then mixed in a 0.15:0.85 (MPAO1:mutant) volumetric ratio, diluted to OD600 0.01 in 3 mL of CAA with and without gentamicin supplementation in a 50 mL falcon tube and grown at 37°C, 200 rpm for 48 h. Samples were taken at 0, 18, 24 and 48 h and diluted in PBS prior to flow cytometry. Data was analysed using a Attune NxT cytometer, counting 5·10^4^ single events per sample. Data analysis was performed using FlowJo V10 (BD Biosciences, Europe). Flow cytometer settings, gating and representative workflow can be found in supplementary methods. Supernatants were collected to analyse gentamicin degradation using LC-MS.

To study competition in biofilms, 30 µL of OD600 0.01 cells were inoculated in a µ-Slide VI – Flat (Ibidi, Germany) and incubated at 37°C in a humid chamber with no agitation for 48 hours. Confocal images (Nikon A1M) were collected from 10 random areas with a 60x Plan Apo lens. A single argon laser line was used with excitation wavelength of 488 nm and using an emission Nikon-FITC filter (525/50 nm) for the imaging of GFP-tagged bacteria; for RFP cells, the excitation wavelength used was 561 nm with the Nikon-mCherry (595/50 nm) emission filter. Images were quantified using colour pixel counter plugin from ImageJ.

### Liquid chromatography and mass spectrometry

We used mass spectrometry determination to confirm the stability of gentamicin over the course of the competition experiments. Five replicates of 5 *µ*L for each sample and standard were injected into an Ultimate 3000 UHPLC ™ system (Thermo Scientific, Bremen, Germany). The analytes were separated using a Kinetex C18 column (100 × 2.1 mm, 5 µm) at a flow rate of 0.25 mL·min^-1^. The autosampler temperature was set to 4°C and the column temperature to 30°C. The initial mobile phase was 99% H_2_O and 1% ACN (both with 0.1% FA) which was increased to 80% ACN and 20% H_2_O (both with 0.1% FA) over 2 minutes and then kept constant for 30 seconds. Subsequently, the mobile phase reverted to the initial composition and was allowed to equilibrate for 30 seconds before commencing the next injection. The total run time was 3 minutes. The UHPLC system was coupled to a ThermoFisher Scientific Q Exactive™ Plus Hybrid Quadrupole mass spectrometer. The electrospray ionization source was operated with a spray voltage of 4.25 kV and a capillary temperature of 390°C. Data was acquired in positive mode, at a mass range of *m/z* 75 to 1,000 with a resolving power of 70,000 (at *m/z* 200), and with the automatic gain control (AGC) on and set to 10^6^ ions. Concentrations were determined using peak ion counts for the gentamicin C1 species (*m/z* 478.3240 ± 5 ppm) measured relative to a known standard concentration of gentamicin of 20.0 µg·mL^-1^. The gentamicin C1 species gave the highest intensity of signal (compared with the gentamicin C1a and C2 / C2a species), with an elution time of 50 seconds. The relationship between peak ion count and concentration was validated through the construction of a calibration curve using standards ranging from 0.1 *µ*g·mL^-1^ to 20.0 *µ*g·mL^-1^.

### *In vivo* studies

*Galleria mellonella* larvae were acquired from a local supplier. Larvae were stored at 4°C and used within two weeks. As before, once cells reached mid-exponential phase, they were washed 3 times in PBS, and the OD600 was normalized to 1. Cells were then mixed in a 0.5:0.5 (MPAO1:mutant) volumetric ratio. Prior to infection, larvae were kept on ice for half an hour to prevent spontaneous movements and then dipped in 70% ethanol to remove any contaminants present on the tegument. Cells were further diluted so that a 10 µL injection in *G. mellonella* would contain bacteria in the range of 10^2^-10^3^ cells. Following this, a stock solution of gentamicin was diluted accordingly in PBS to obtain a 20 µg·mL^-1^ final concentration in the larvae after a 10 µL injection, assuming that every larvae as a volume of 1 mL. Both bacteria and gentamicin were injected in the last pair of prolegs using a Hamilton syringe and a peristaltic pump. Syringes were discarded after each treatment to prevent carryover between samples. Larvae were incubated at 37°C and their survival was checked every hour for mono-infections. Larvae were considered dead if there was no sudden movement after touch with a pipette tip on the head and culled at 15 hours with the haemocoel being recovered. To do so, larvae were kept in ice for 30 minutes until no movement was detected, then a small incision was generated using a sterile scalpel to recover haemocoel, which was diluted and 20 µL were plated on Pseudomonas Isolation Agar supplemented with gentamicin to calculate population proportions. Cells were allowed to grow for 18 hours at 37°C and then stored for 3 days at 4°C, which facilitates their differentiation through fluorescent reporter accumulation in the bacterial colonies.

### Data analyses

All graphs were plotted using ggplot2 (R package). OD was fitted using Fitderiv (43) in order to obtain derivative curves accounting for growth rates at each time point. Pyoverdine production per cell was calculated as the fluorescence at 450 nm(Ex)/490 nm(Em) divided by OD600 at each time point. The area under the curve (AUC) of pyoverdine production up to the maximum growth rate was obtained using the OD600 derivative curve calculated with Fitderiv as an indicator of the point of maximum growth rate. The calculations of AUC were performed using Graphpad Prism 7.

All experiments were conducted from the beginning on three different days (biological replicates) carrying at least three technical replicates each time. One-way ANOVA complemented with Dunnett post-hoc tests were used to analyse the significance of differences found in growth rates, pyoverdine production up to the maximum growth rate and the frequency of strains observed in biofilm formation experiments. The control group for comparison in each assay was MPAO1 in the corresponding condition, except in the biofilm experiments, in which samples were compared to the frequencies obtained in the MPAO-Δ*pvdA* (-Gm) competition. t-tests were used for the analysis of growth rates and pyoverdine production up to maximum growth rate when overexpressing *fpvB*, as well as in the competition studies in unstructured environments and in *G. mellonella*. The control groups were the corresponding ‘empty’ construct in *fpvB* overexpression experiments or the initial frequencies of each of the strains in the competitions. The survival curve analyses were performed using Long-Rank test from survminer, an R package. In all cases, the level of significance was established at p ≤ 0.05, p ≤ 0.01 and p ≤ 0.001 represented in the figures with, respectively, one, two or three asterisks.

## Acknowledgements

The authors would like to thank Dr Melanie Ghoul, Prof. Stuart West and Prof. Craig McLean (University of Oxford), and Dr Jorge Gutiérrez (University of Surrey) for the insightful discussions on population invasion and their constructive feedback on the manuscript. The authors are indebted to Dr Helen King, Dr Mandy Fivian-Hughes, Kate Reid and Anita Sicilia for their technical assistance. JG was the recipient of a PhD studentship of the University of Surrey and an EMBO Short-Term Fellowship (ASFT number: 8166). CAR and JJ acknowledge the support received from the Biotechnology and Biological Sciences Research Council (BBSRC) (grants BB/L02683X/1 and BB/T011289/1 from the ERA-Cobiotech programme of the EU). MS, CC and MB acknowledge the support from the Engineering and Physical Sciences Research Council through a strategic equipment grant (EP/P001440/1). RK has received funding from the European Research Council (ERC) under the European Union’s Horizon 2020 research and innovation programme (grant agreement no. 681295). ÖÖ has received funding from the Forschungskredit by the University of Zurich.

## Author contributions

JG, CAR, RK and JJ designed and supervised the study. JG, MS1, ÖÖ, MS2 and CC conducted experimental work. All authors analysed the data and wrote the manuscript.

