## Supplementary information for "The loss of the pyoverdine secondary receptor in *Pseudomonas aeruginosa* results in a fitter strain suitable for population invasion"

### Supplementary results

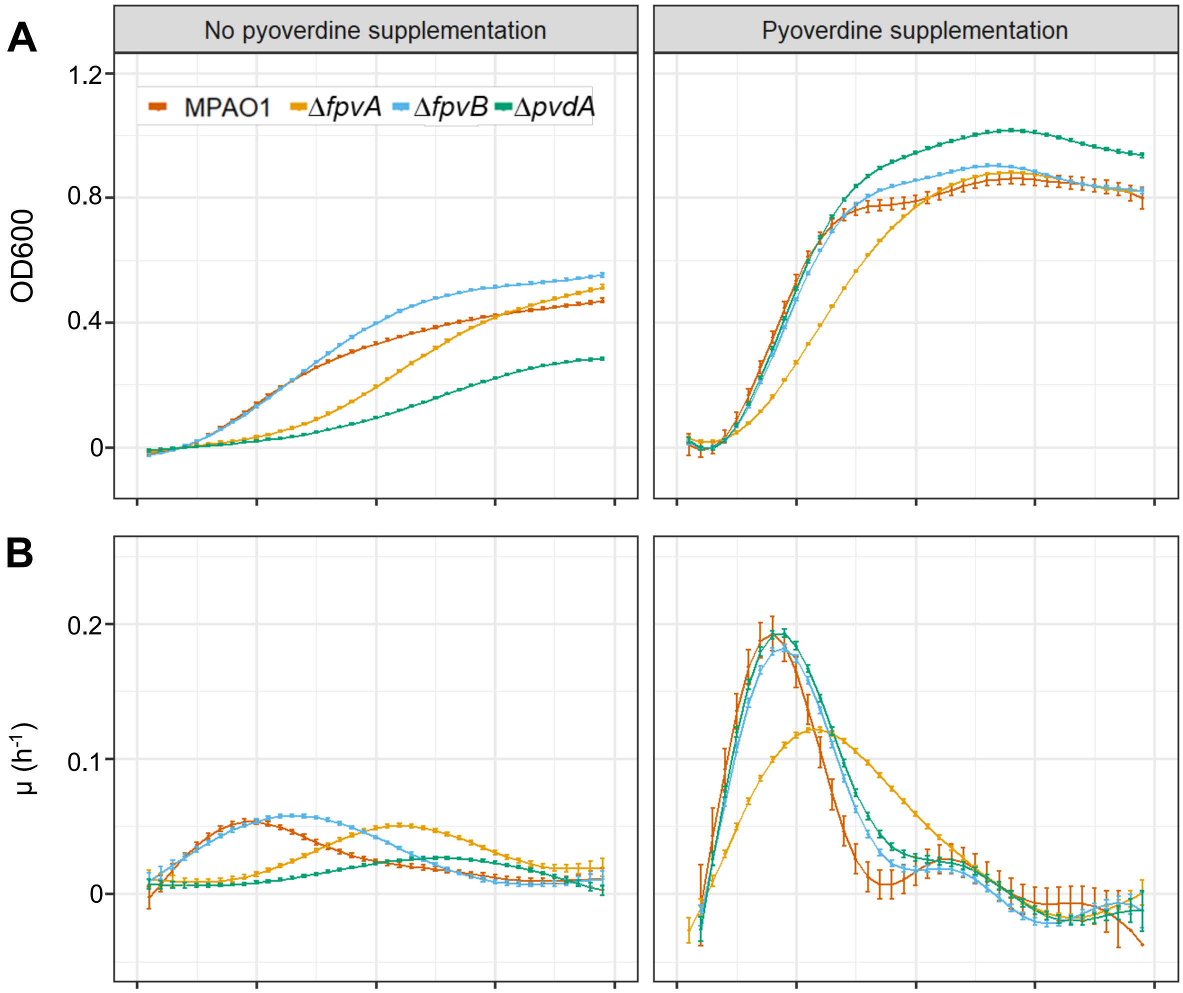


**Fig. S1. Kinetics of bacterial growth and pyoverdine production.** (A) Growth kinetics of MPAO1 and mutants in iron-limited CAA in the absence (left) and presence (right) of exogenous pyoverdine. (B) First derivative of growth curves obtained with fitderiv (1). Error bars represent standard error of the mean from three biological replicates.

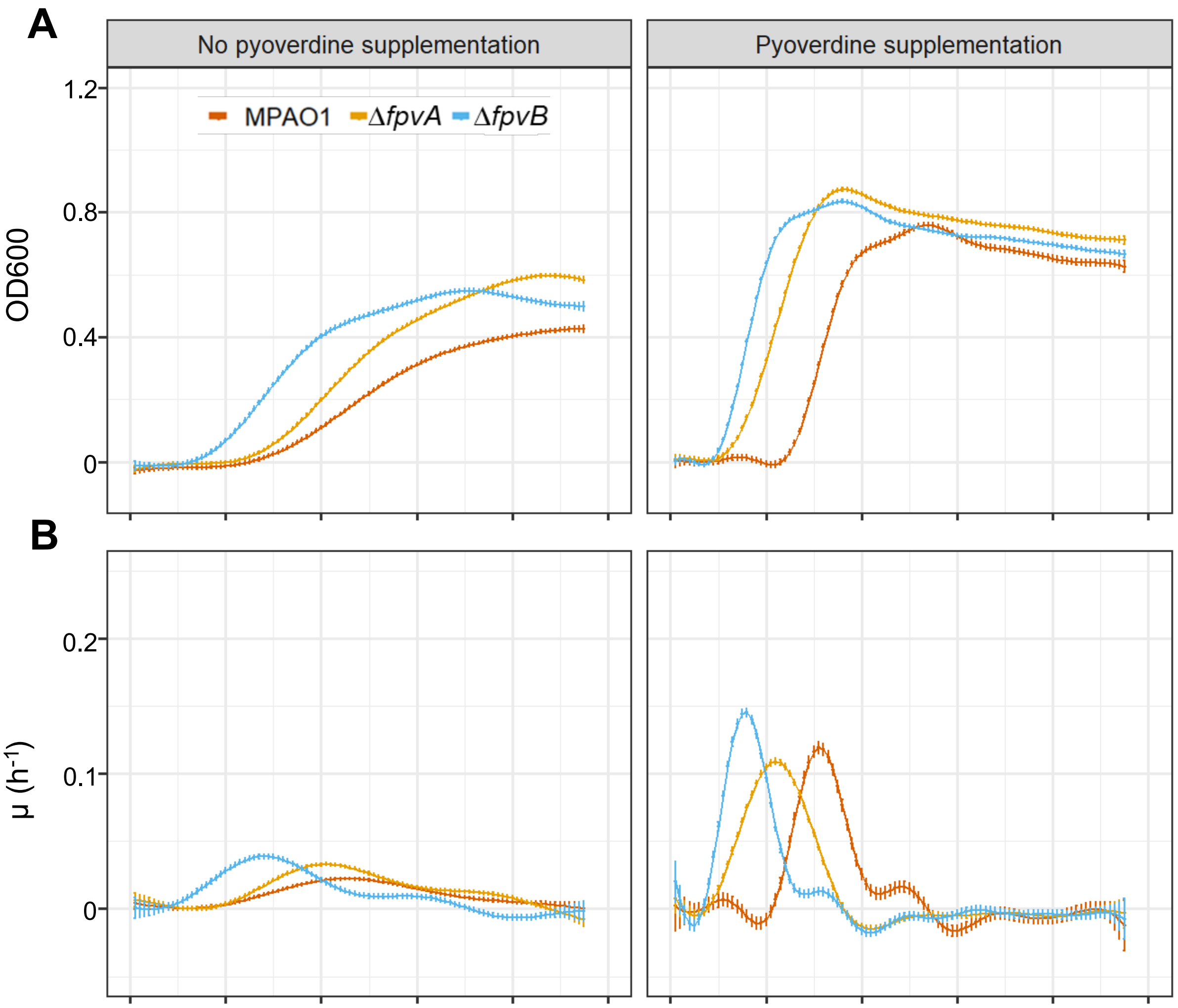


**Fig. S2. Kinetics of bacterial growth and pyoverdine production in the presence of gentamicin.** (A) Growth kinetics of MPAO1 and mutants in iron-limited CAA in the absence (left) and presence (right) of exogenous pyoverdine. (B) First derivative of growth curves obtained with fitderiv (1). Error bars represent standard error of the mean from three biological replicates.

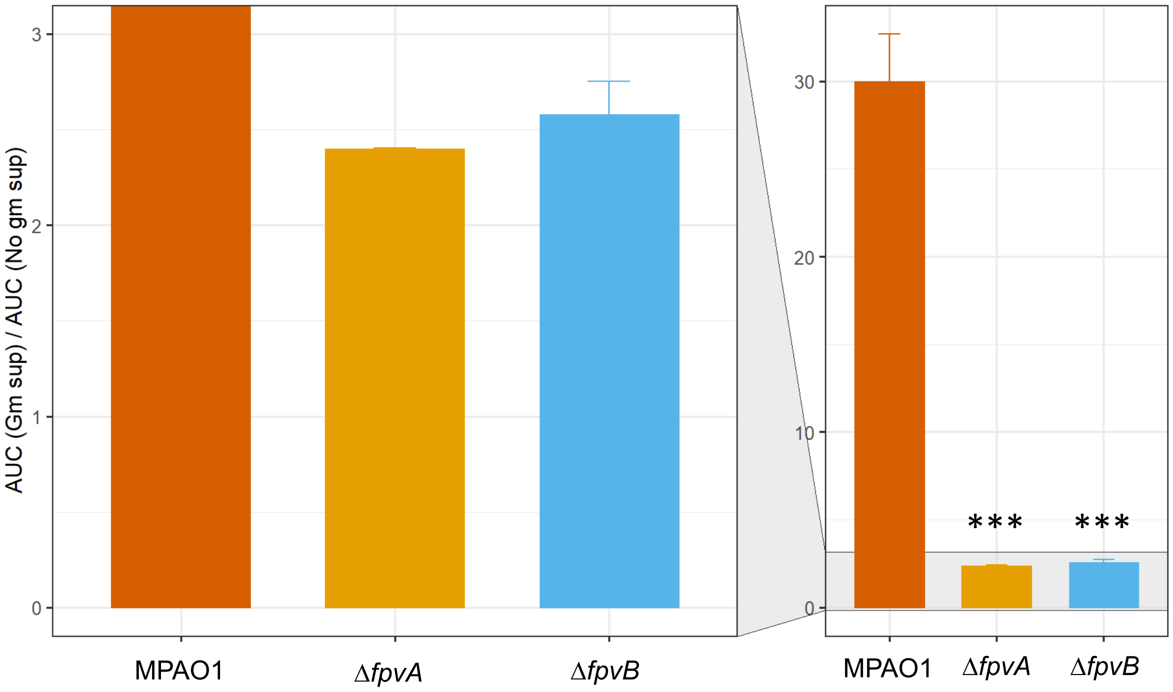


**Fig. S3. Analysis of the influence of gentamicin on pyoverdine production up to maximum growth rate in iron limited CAA without pyoverdine supplementation.** The bars represent the ratio between the pyoverdine production per cell up to the maximum growth rate divided by in the corresponding value obtained in the absence of the antibiotic. All ratios are represented in the right panel while the left panel zooms in the lower range to allow comparison between the Δ*fpvA* and Δ*fpvB* ratios. Error bars, corrected for error propagation, represent the standard error of the mean from three biological replicates.

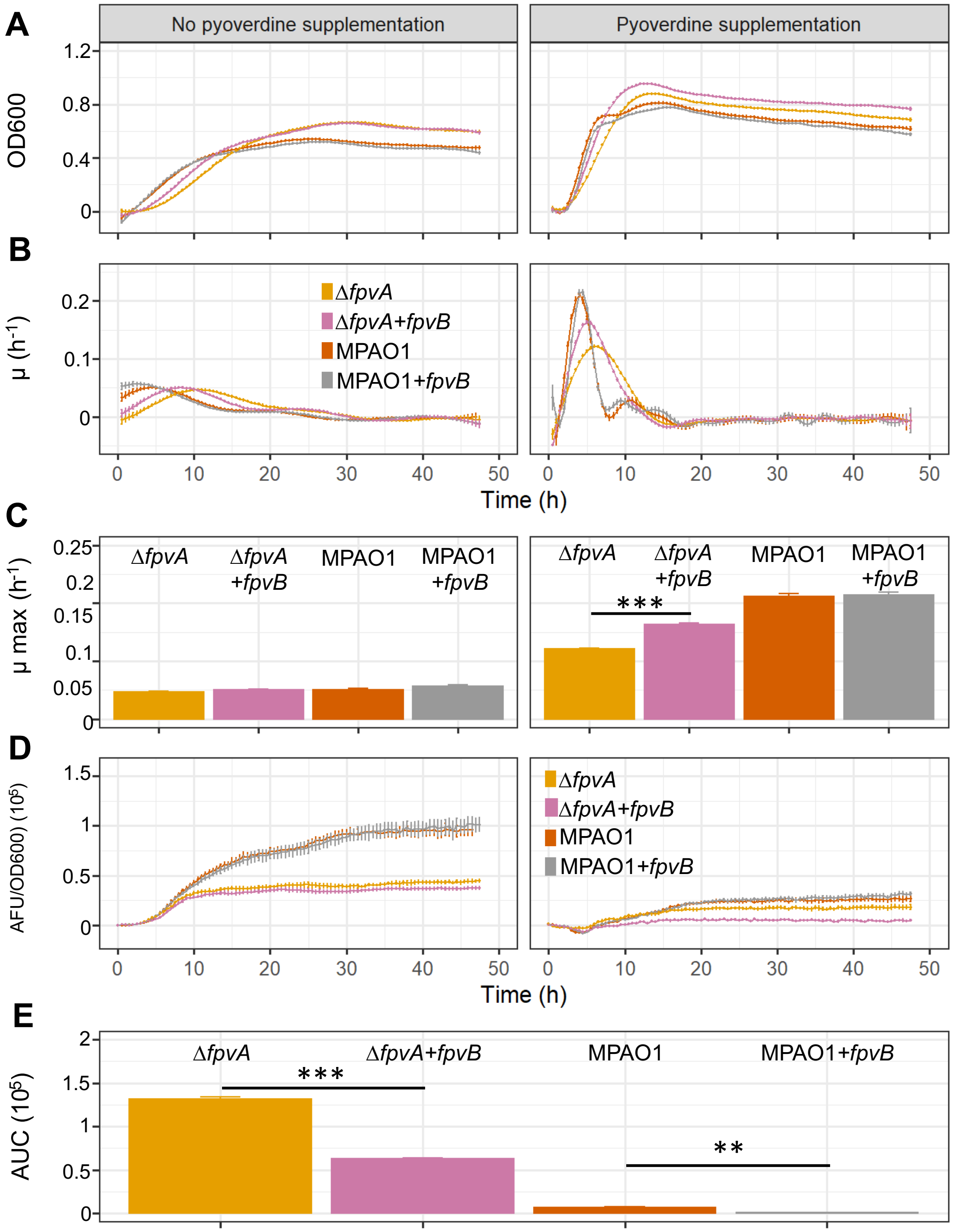


**Fig. S4. Effect of an extra copy of *fpvB* in the kinetics of bacterial growth and pyoverdine production in the absence of gentamicin.** (A) Growth kinetics of MPAO1 and Δ*fpvA* with and without an extra copy of *fpvB* in iron-limited CAA in the absence (left) and presence (right) of exogenous pyoverdine. (B) First derivative of growth curves obtained with fitderiv (1). (C) Maximum growth rates. (D) Pyoverdine production per cell over time of the cultures in same conditions. (E) Pyoverdine accumulated in cultures of each of the strains until reaching the maximum growth rate. without exogenous pyoverdine. Error bars represent standard error of the mean from three biological replicates. Asterisks show significant differences compared to MPAO1 in a One-way ANOVA test. Expression of FpvB did not affect the growth profile of MPAO1 but yielded a distinguishable advantage in Δ*fpvA* where it also significantly reduced the amount of pyoverdine required to reach the maximum growth rate. This observation is consistent with FpvB diverting ferripyoverdine from the FpvA via of entry to the cells and thus interfering with the autocatalytic activation of pyoverdine synthesis in this regulatory pathway.

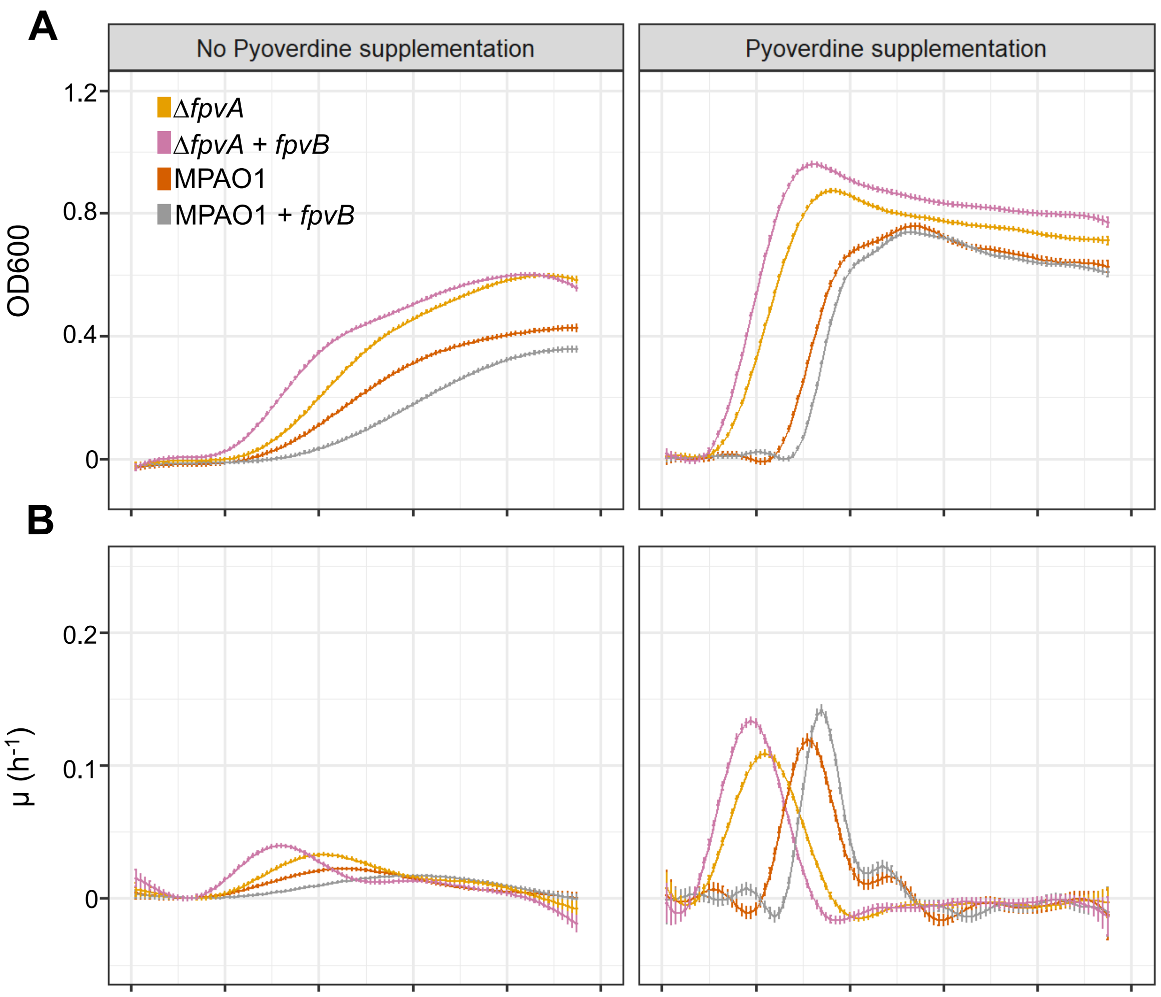


**Fig. S5. Effect of an extra copy of *fpvB* in the kinetics of bacterial growth and pyoverdine production in the presence of gentamicin.** (A) Growth kinetics of MPAO1 and Δ*fpvA* with and without an extra copy of *fpvB* in iron-limited CAA in the absence (left) and presence (right) of exogenous pyoverdine. (B) First derivative of growth curves obtained with fitderiv (1). Error bars represent standard error of the mean from three biological replicates.

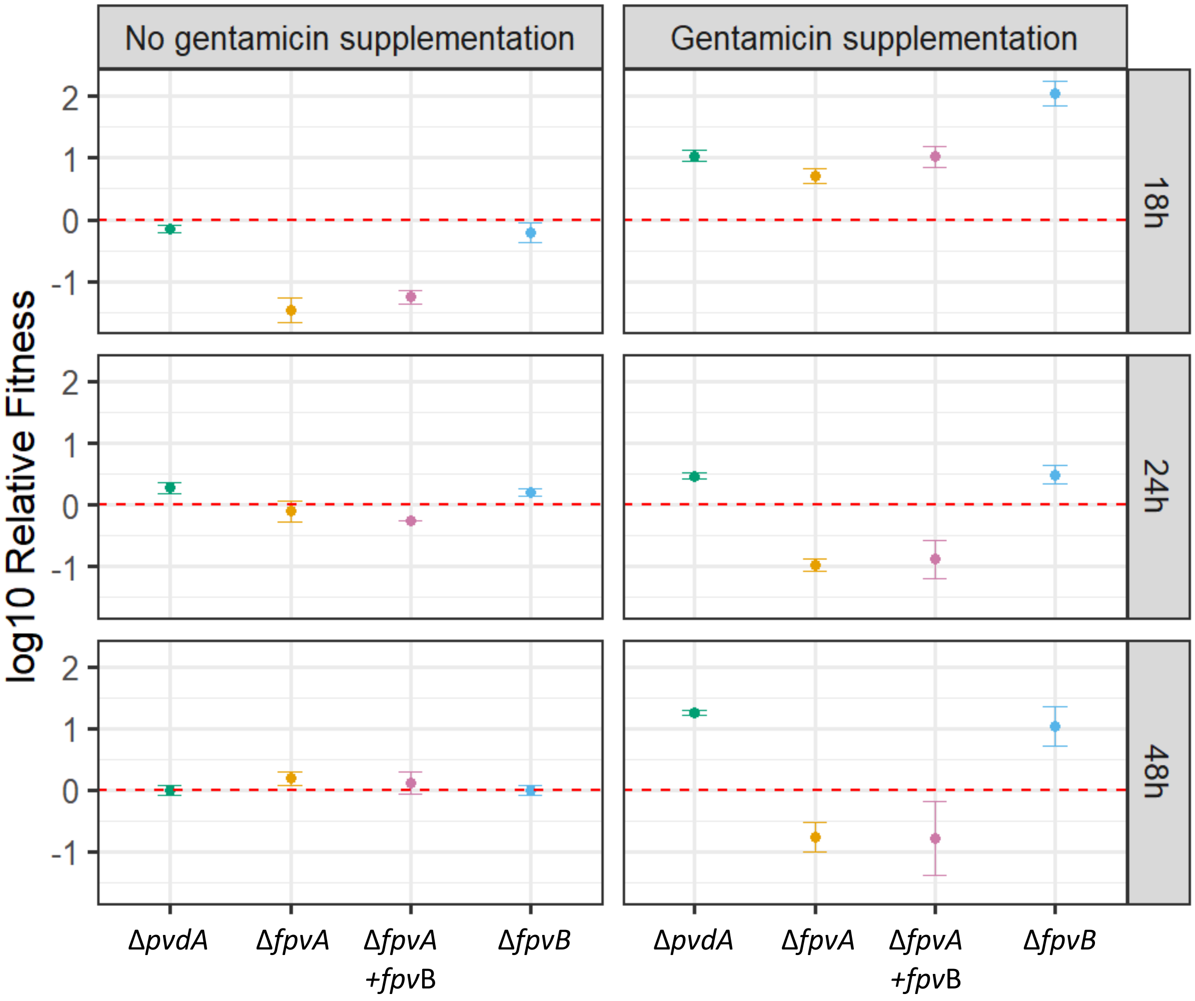


**Fig. S6. Relative fitness of the mutant strains compared to MPAO1 in iron limited CAA in the presence or absence of gentamicin.** The log10 relative fitness (v) of mutants was calculated as v = [log10($t_2$)(1− log10($t_1$))]/[log10($t_1$)(1− log10($t_2$))], where $t_1$ is the initial proportion of mutants and $t_2$ is their final proportion. The fitness value of v therefore signifies whether mutants increased in (v > 0), decreased (v < 0) or remained at the same frequency (v = 0) over the duration of the competition assay. Error bars represent CI 95% of the mean from three biological replicates with alternate fluorescent tags (GFP and RFP).

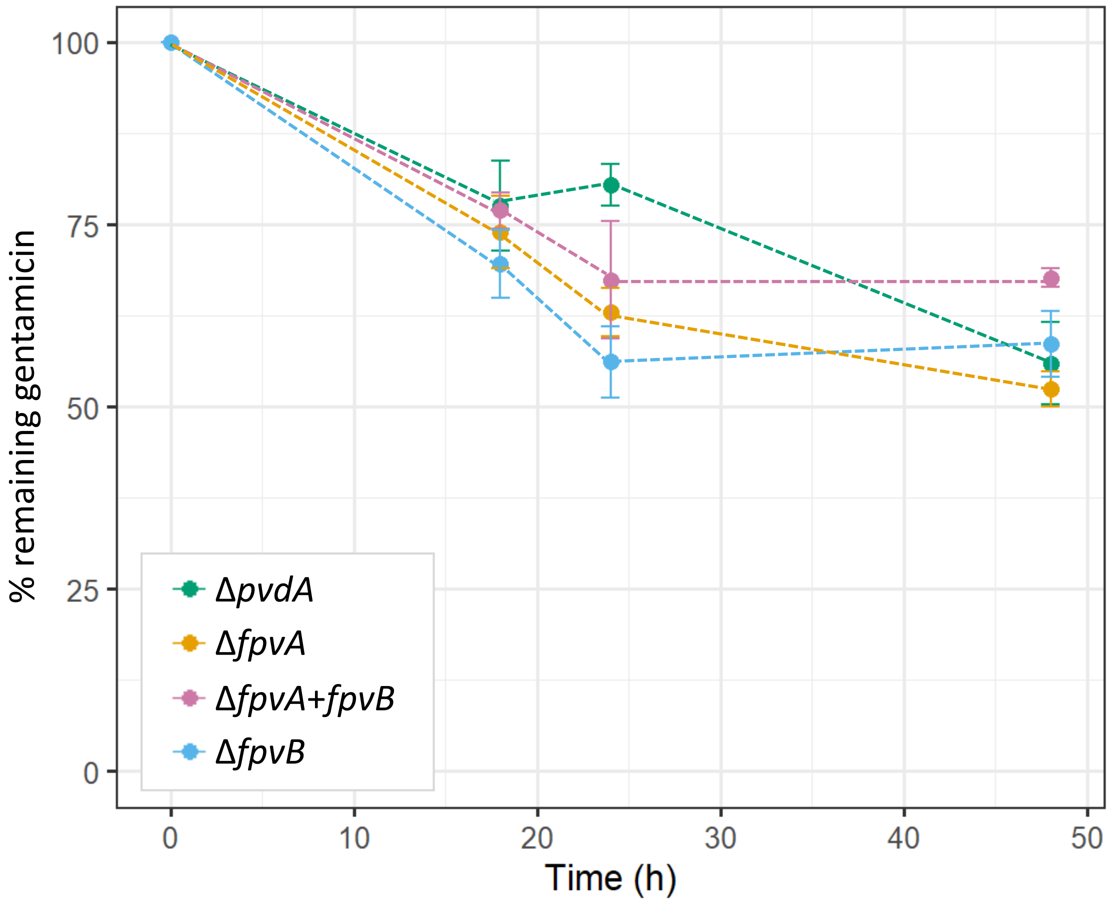


**Fig. S7. Kinetics of gentamicin degradation in competition assays between MPAO1 and different mutants.** Aliquots of culture supernatants were taken at different times and analysed by mass spectrometry. Results correspond to the mean and standard deviation of two biological replicates of each time point measured five times. The results showed that half of the gentamicin remained in the cultures by the end of the experiment and only 20-35% of the antibiotic is degraded at 18 h when MPAO1 starts increasing its relative fitness. The final concentration of gentamicin found (10 $\mu g \cdot mL^{-1}$) is 5 times higher than the minimum inhibitory concentration reported for MPAO1 (2).

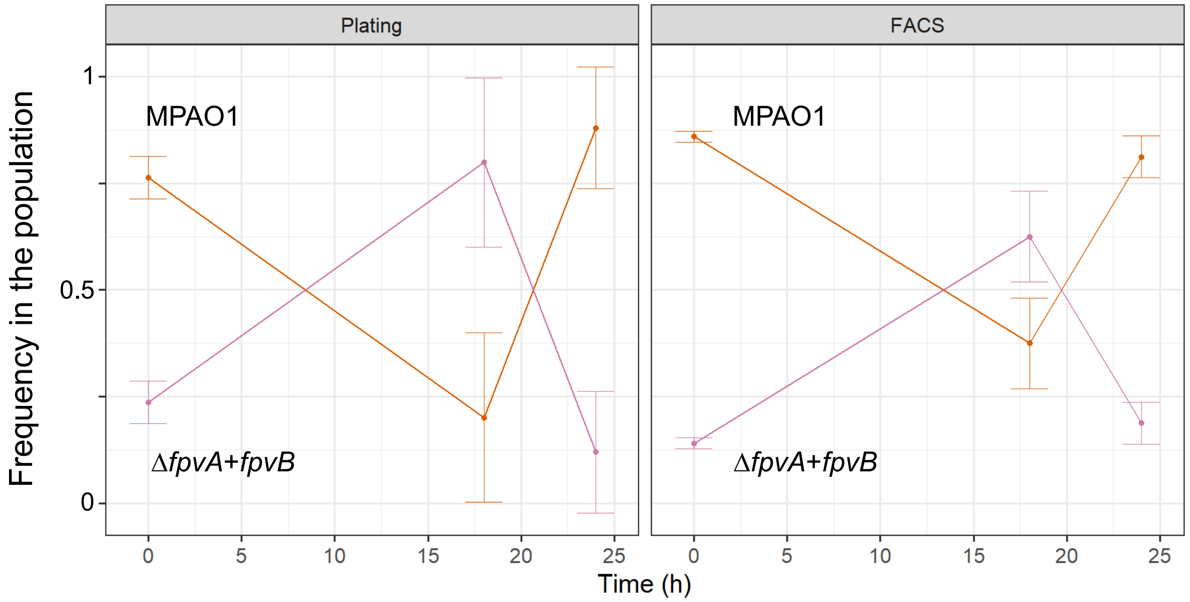


**Fig. S8. Comparison between cfu determination (plating) and flow cytometry (FACS) for population dynamics monitoring.** The results correspond to competition assays between the strains indicated in iron limited CAA supplemented with gentamicin. Bacterial proportions were calculated either based on $5 \cdot 10^4$ event counts in a flow cytometer or after plating on LB plates following serial dilutions. Error bars represent standard error of the mean from three biological replicates with alternate fluorescent tags (GFP and RFP).

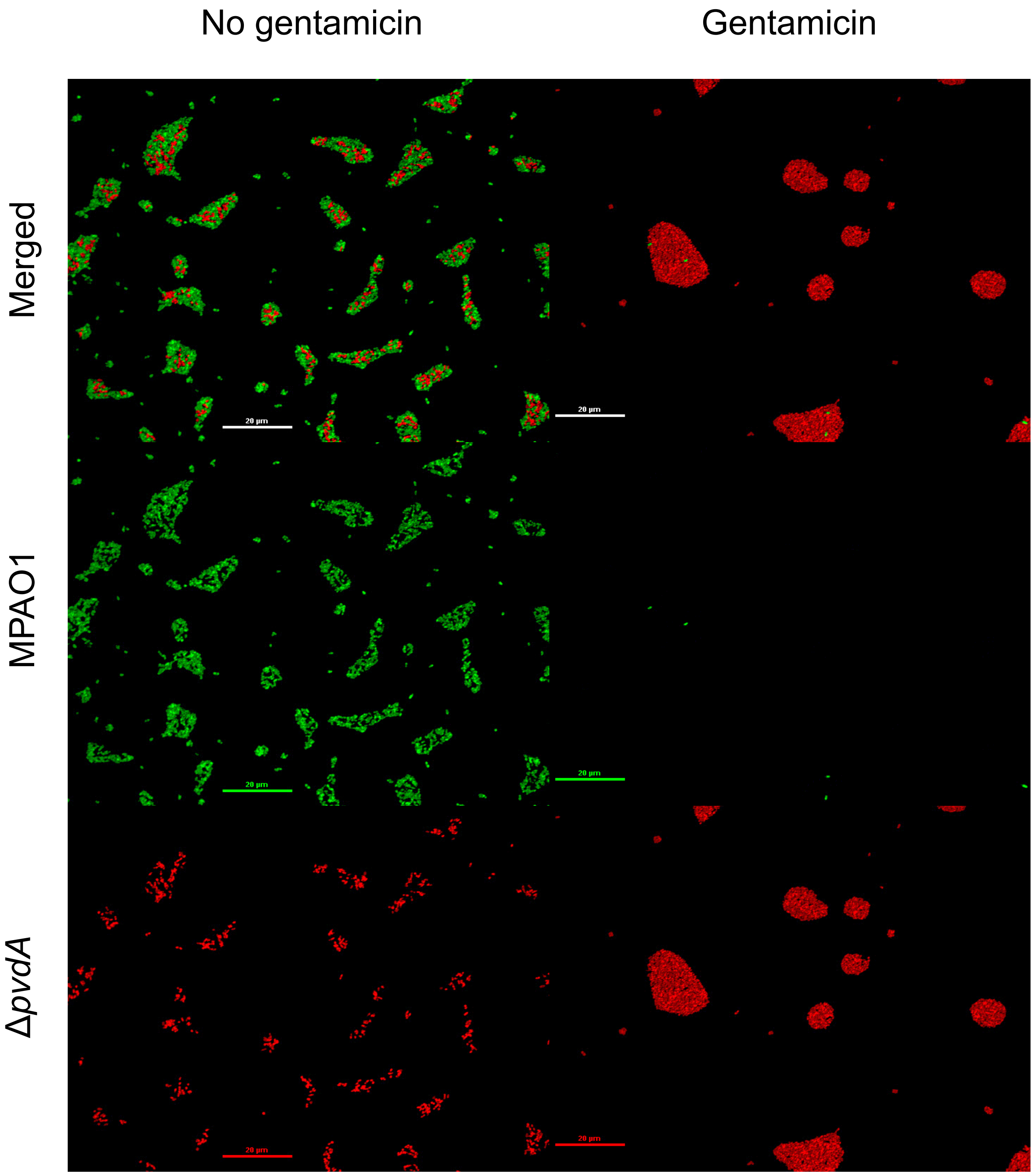


**Fig. S9.** ***In vitro* biofilm competition studies between MPAO1 and Δ*pvdA*.** After 48 hours growing in iron limited CAA, cells were imaged using a Nikon A1M confocal microscope. MPAO1 was tagged using GFP and Δ*pvdA* using RFP. Each image is representative of a set of 10 randomly selected areas (selected using phase contrast to avoid bias) for each condition, taken for each biological replicate. Scale bars equal 20 $\mu$m.

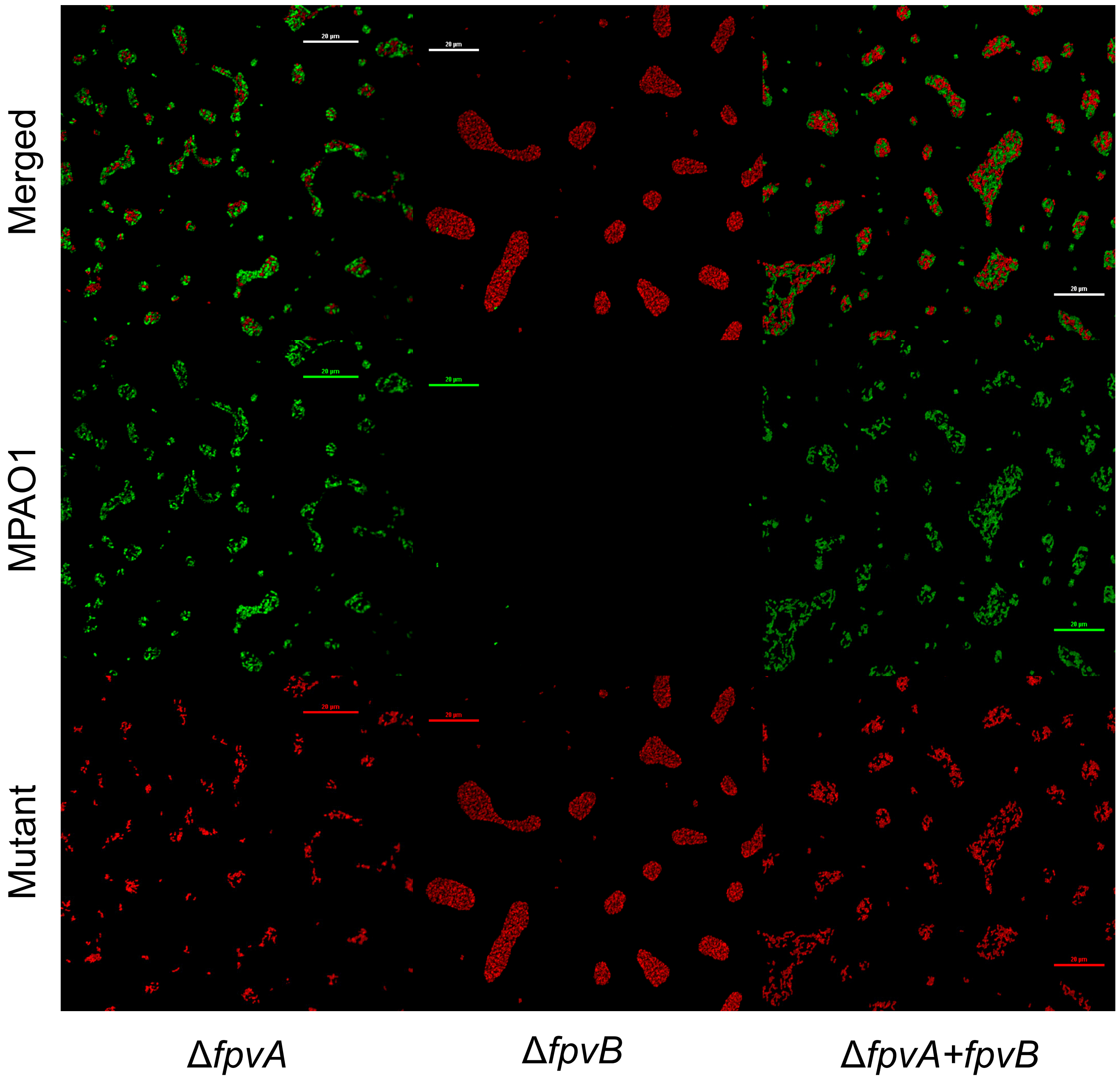


**Fig. S10. *In vitro* biofilm competition studies between MPAO1 and the mutants Δ*fpvA*, Δ*fpvB and* Δ*fpvA*+*fpvB*.** After 48 hours growing in iron limited CAA, cells were imaged using a Nikon A1M confocal microscope. MPAO1 was tagged using GFP and Δ*pvdA* using RFP. Each image is representative of a set of 10 randomly selected areas (selected using phase contrast to avoid bias) for each condition, taken for each biological replicate. Scale bars equal 20 $\mu$m.

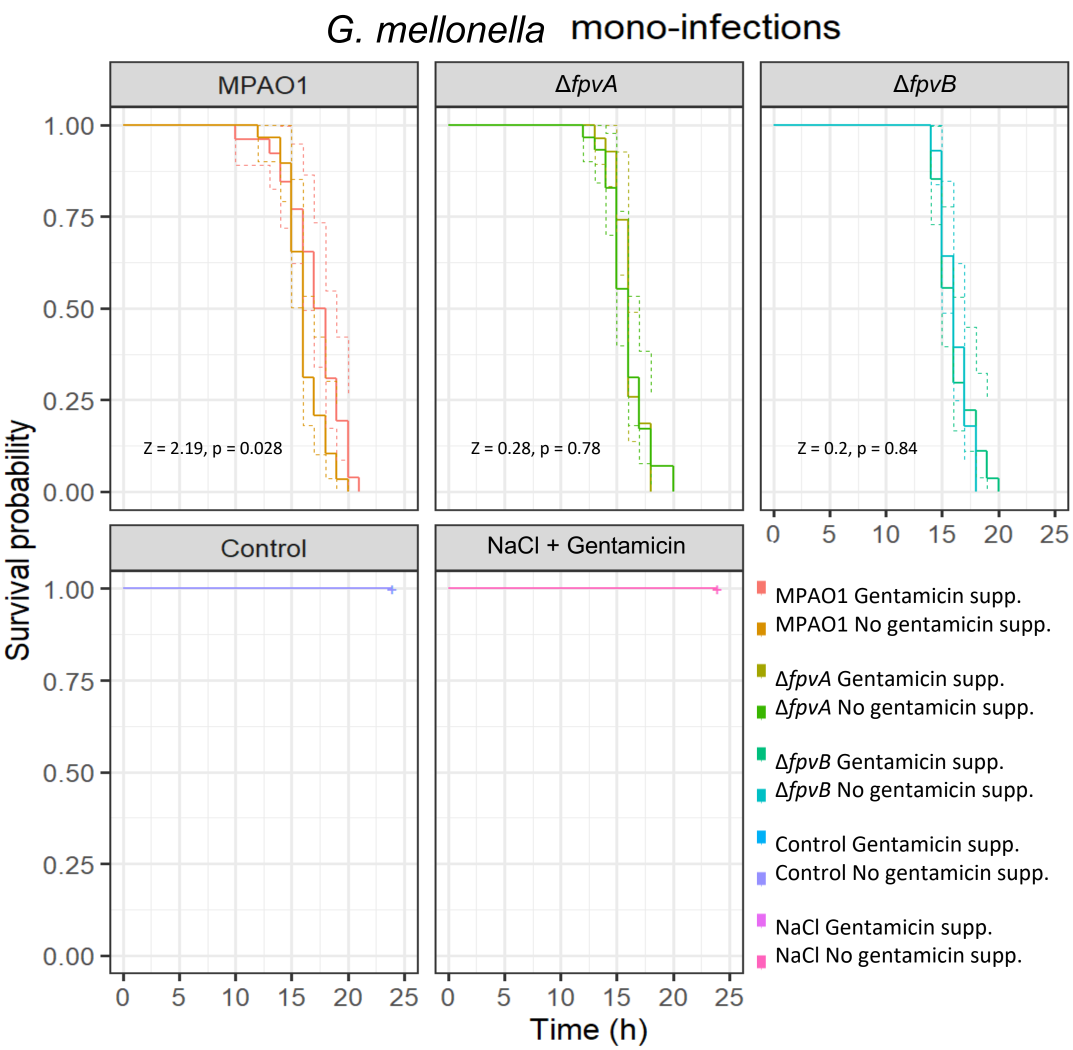


**Fig. S11. Survival of *G. mellonella* during mono-infections**. The survival of the larvae was monitored every hour after initial infection. Individuals not responding to physical stimulus were considered dead, independently of the degree of melanisation. Lines represent the mean value of three biological replicates. Dotted lines represent confidence intervals (95%) obtained in a Long-Rank test.

### Supplementary methods

All strains, plasmids and primers used in this study are listed, respectively, in tables S1, S2 and S3.

#### *Construction of fpvA* and *fpvB* promoter fusions to a fluorescent reporter

A graphical summary of the cloning strategy for all plasmids is depicted in Fig. S12. Initially, msfGFP was replaced with mCherry in pBG25 to avoid possible signal overlaps between pyoverdine and the fluorescent reporter protein. To do so, mCherry was amplified using the mChFWeco and mChREVbam oligonucleotides containing EcoRI/BamHI sites. The PCR product together with pBG25 were digested with EcoRI/BamHI followed by ligation and transformation in One Shot™ PIR2 *E. coli*. The primers used to test whether the ligation process was successful were mChFWconf and mChREVconf. The resulting plasmid, pBG25-mCherry, contained the 14F promoter (a strong synthetically engineered promoter), a bicistronic linker carrying a secondary ribosome binding site (RBS) and mCherry.

In the next step, the promoter regions of *fpvA* and *fpvB* genes were cloned into pBG25-mCherry. The bicistronic linker present in the pBGXX plasmids was introduced to solve the problem of translational variability caused by mRNA secondary structure, thus making promoter activity comparable between genes by normalizing translation (3,4). mRNA secondary structure in the region between -4 and +37 relative to the translation star site accounts for almost 60% of the protein expression variability (5). pBGXX plasmids contain a 5´UTR with two ribosome binding sites: the first one initiates translation of a short peptide while the second one is integrated in the sequence of this short peptide and defines the rate of translation of the downstream gene of interest (3,4). This short peptide (with low mRNA secondary structure) contains a termination codon overlapped to the start codon of the gene of interest, so the ribosome does not detach from the mRNA when the short peptide translation is finished (4). By including this motif in all constructs it is possible to account for the majority of the gene expression variability and further changes can be attributed to changes in the promoters (4).

The *fpvA* promoter of *P. aeruginosa* MPAO1 was obtained as a pair of complementary oligonucleotides (fpvApFWpac and fpvApREVavr) containing the whole sequence and suitable restriction sites. The oligonucleotides were annealed using T4 DNA ligase buffer (without the enzyme) heated to 95ºC for five minutes in a water bath and allowed to cool down to room temperature. The *fpvB* promoter region of *P. aeruginosa* MPAO1 was PCR amplified using fpvBpFWpac and fpvBpREVavr primers**.** Both promoters (*fpvA* and *fpvB*) and pBG25-mCherry were digested using PacI /AvrII, ligated and transformed, resulting in pBG25-fpvAp-mCherry and pBG25-fpvBp-mCherry. The primers used to test whether the ligation process was successful were mChFWconf and mChREVconf. Constructions were then integrated into the Tn7 site of the different *P. aeruginosa* strains following the protocol described in the methods section of the main text. Confirmation of the correct insertion of the desired fragments was done by PCR using both Tn7RFW and glmsDownREV primers, followed by DNA Sanger sequencing (Genewiz, UK).

#### *Construction of gentamicin resistant strains*

A control plasmid was constructed by digesting pBG25-mCherry with PacI/XbaI, followed by blunting (Quick Blunt Kit -NEB-) and religation, generating the plasmid pBG-control. Integration into Tn7 locations generated *P. aeruginosa* MPAO1::Tn7*:aacC1, P. aeruginosa* Δ*fpvA::*Tn7*:aacC1*, Δ*fpvB::*Tn7*:aacC1 and* Δ*pvdA::*Tn7*:aacC1* strains, all resistant to gentamicin carrying no expression systems. The correct modification of the strains was confirmed by PCR and Sanger sequencing (Genewiz, UK)

#### *Construction of strains with an extra copy of the fpvB secondary receptor*

In order to obtain strains able to overexpress *fpvB,* the natural RBS and the gene *fpvB* from *P. aeruginosa* MPAO1 were PCR amplified with the corresponding restriction sites (XbaI/HindIII) added at the 5´ end of the primers fpvB+RBSFWxba and fpvB+RBSREVhind. pBG25 was digested using AvrII/HindIII (note that XbaI and AvrII present compatible ends). Both fragments, pBG25 backbone and the *fpvB* gene were then ligated and transformed into in One Shot™ PIR2 *E. coli*. The primers used to test whether the ligation process was successful were fpvB+RBSFWconf and fpvB+RBSREVconf. The plasmid generated, pBG25-14FfpvB (Fig. S12), carried the construction that was integrated in the *fpvB* locus of the chromosome of *P. aeruginosa* MPAO1 and *P. aeruginosa* Δ*fpvA* following the protocol described in the main text, generating *P. aeruginosa* MPAO1::*aacC1:14FfpvB and P. aeruginosa* Δ*fpvA*::*aacC1*:*14FfpvB*. The resulting strains harbour two functional copies of the *fpvB* gene, one expressed heterologously from the 14F promoter. The correct modification of the strains was confirmed by whole genome sequencing (MicrobesNG, UK).

***Construction of fluorescently tagged strains for competition studies***
In order to allow for proper comparisons between strains, all competition experiments were conducted with a modified version of the different mutants that reproduce the genetic architecture of the Tn7 insertion in the engineered *P. aeruginosa ΔfpvA::*Tn7*:aacC1:14FfpvB*. Strains were constructed so they would share the same DNA insert with the exception of the *fpvB* gene. These strains were generated as follows: pBG25-mCherry was digested with EcoRI and religated, obtaining pBG25-control14F. In addition, several of the competing strains were pyoverdine producers and they were tagged with fluorescent reporters to allow their differentiation.

The promoter *Ptet* for the constitutively expression of fluorescent tags and the corresponding transcriptional terminators were amplified from MBP-1.0 and MBP-swap (6) using two different sets of primers: GFP/RFPFW_MBPpac together with GFP/RFPREV_MBPxho and GFP/RFPFW_MBPpac with GFP/RFPREV_MBPbam, containing PacI/XhoI and PacI/BamHI restriction sites, respectively, for each of the destination backbones. Equally, backbone plasmids were amplified using BG25_C14FFWpac with BG25_C14FREVxho and BG25_14FBFWpac with BG25_C14FBREVbam primers containing the corresponding specific restriction sites (PacI/XhoI and PacI/BamHI). Therefore, when cloning into pBG25-control14F, PacI/XhoI were used as restriction enzymes. When pBG25-14FfpvB was the recipient backbone, PacI/BamHI were used. The ligation of the corresponding fragments yielded 2 plasmids with alternate colours (4 in total): pBG25-GFP/RPF-control14F and pBG25-GFP/RFP-14FfpvB (Figs. S13). The primers used to test whether the ligation process was successful were BG25_TAGFW, BG25_TAG_GFPREV and BG25_TAG_RFPREV. To remove the unused 14F promoter, pBG25-GFP/RPF-control14F was digested with PacI/XbaI, followed by blunting (Quick Blunt Kit -NEB-) and religation, generating the plasmids pBG25-GFP/RPF-control. The transposons carrying the required DNA inserts were delivered by triparental mating yielding: *P. aeruginosa* MPAO1::Tn7:*aacC1*:GFP/RFP, *P. aeruginosa* Δ*fpvA*::Tn7:*aacC1*:GFP/RFP, *P. aeruginosa* Δ*fpvB*::Tn7:*aacC1*:GFP/RFP, *P. aeruginosa* Δ*pvdA*::Tn7:*aacC1*:GFP/RFP and *P. aeruginosa* Δ*fpvA*::*aacC1*:GFP/RFP:*14FfpvB*. Insertions were confirmed by PCR and sequencing (Genewiz, UK).

**Table S1.** List of bacterial strains used in this study

| Strain | Description | Source |
|---|---|---|
| *P. aeruginosa* MPAO1 | Parental strain/Wildtype | (7) |
| *P. aeruginosa* Δ*pvdA* | Defective strain for pyoverdine synthesis. PW5012. *P. aeruginosa* gene no. PA2386, gene product: L-ornithine N5-oxygenase. Transposon ISlacZ/hah. | (7) |
| *P. aeruginosa* Δ*fpvA* | Defective strain for pyoverdine primary receptor. PW5036. *P. aeruginosa* gene no. PA2398, gene product: FpvA pyoverdine receptor.Transposon ISlacZ/hah. | (7) |
| *P. aeruginosa* Δ*fpvB* | Defective strain for pyoverdine secondary receptor. PW8065. *P. aeruginosa* gene no. PA4168, gene product: FpvB pyoverdine receptor.Transposon ISlacZ/hah. | (7) |
| *P. aeruginosa* Δ*pchE* | Defective strain for pyochelin receptor. | (7) |

| | | |
|---|---|---|
| | PW8174. *P. aeruginosa* gene no. PA4226, gene product: dihydroaeruginoic acid synthetase.Transposon ISpho/hah. | |
| *P. aeruginosa* MPAO1::Tn7:*aacC1* | *P. aeruginosa* MPAO1 carrying *aacC1*, conferring gentamicin resistance in the Tn7 location | This study |
| *P. aeruginosa* Δ*pvdA*::Tn7:*aacC1* | *P. aeruginosa* Δ*pvdA* carrying *aacC1*, conferring gentamicin resistance in the Tn7 location | This study |
| *P. aeruginosa* Δ*fpvA*::Tn7:*aacC1* | *P. aeruginosa* Δ*fpvA* carrying *aacC1*, conferring gentamicin resistance in the Tn7 location | This study |
| *P. aeruginosa* Δ*fpvB*::Tn7:*aacC1* | *P. aeruginosa* Δ*fpvB* carrying *aacC1*, conferring gentamicin resistance in the Tn7 location | This study |
| *P. aeruginosa* MPAO1::Tn7:*aacC1*:*fpvAp*:*mCherry* | *P. aeruginosa* MPAO1 with genomic insertion of Tn7-fpvAp-mCherry in the Tn7 location | This study |
| *P. aeruginosa* Δ*fpvA*::Tn7:*aacC1*:*fpvAp*:*mCherry* | *P. aeruginosa* Δ*fpvA* with genomic insertion of Tn7-fpvAp-mCherry in the Tn7 location | This study |
| *P. aeruginosa* Δ*fpvB*::Tn7:*aacC1*:*fpvAp*:*mCherry* | *P. aeruginosa* MPAO1 with genomic insertion of Tn7-fpvAp -mCherry in the Tn7 location | This study |
| *P. aeruginosa* MPAO1::Tn7:*aacC1*:*fpvBp*:*mCherry* | *P. aeruginosa* MPAO1 with genomic insertion of Tn7-fpvBp-mCherry in the Tn7 location | This study |
| *P. aeruginosa* Δ*fpvA*::Tn7:*aacC1*:*fpvBp*:*mCherry* | *P. aeruginosa* Δ*fpvA* with genomic insertion of Tn7-fpvBp-mCherry in the Tn7 location | This study |
| *P. aeruginosa* Δ*fpvB*::Tn7:*aacC1*:*fpvBp*:*mCherry* | *P. aeruginosa* Δ*fpvB* with genomic insertion of Tn7-fpvB -mCherry in the Tn7 location | This study |
| *P. aeruginosa* MPAO1*::aacC1:14FfpvB* | *P. aeruginosa* MPAO1 with genomic insertion of aacC1-14FfpvB in the *fpvB* locus | This study |
| *P. aeruginosa* Δ*fpvA*::*aacC1:14FfpvB* | *P. aeruginosa* Δ*fpvA* with genomic insertion of aacC1-14FfpvB in the *fpvB* locus | This study |
| *P. aeruginosa* MPAO1::Tn7:aacC1:GFP/RFP | *P. aeruginosa* MPAO1 with genomic insertion Tn7-GFP/RPF-control in the Tn7 location | This study |
| *P. aeruginosa* Δ*fpvA*::*Tn7*:*aacC1*:*GFP/RFP* | *P. aeruginosa* Δ*fpvA* with genomic insertion of Tn7-GFP/RPF-control in the Tn7 location | This study |
| *P. aeruginosa* Δ*fpvB*::*Tn7*:*aacC1*:*GFP/RFP* | *P. aeruginosa* Δ*fpvB* with genomic insertion of Tn7-GFP/RPF-control in the Tn7 location | This study |
| *P. aeruginosa* Δ*pvdA*::*Tn7*:*aacC1*:*GFP/RFP* | *P. aeruginosa* Δ*pvdA* with genomic insertion of Tn7-GFP/RPF-control in the Tn7 location | This study |
| *P. aeruginosa* Δ*fpvA*::*aacC1*:*GFP/RFP*:*14FfpvB* | *P. aeruginosa* Δ*fpvA* with genomic insertion of Tn7-GFP/RFP-14FfpvB in the *fpvB* locus | This study |
| *E. coli* DH5*α* | Competent cells. Genotype: *F– endA1 glnV44 thi-1 recA1 relA1 gyrA96 deoR* | Lab collection |

| | | |
|---|---|---|
| | *nupG purB20 φ80dlacZΔM15 Δ(lacZYA-argF)U169, hsdR17(rK–mK+), λ–.* | |
| One Shot™ PIR2 Chemically Competent *E. coli* | Competent cells. Genotype *F-Δlac169 rpoS(Am) robA1 creC510 hsdR514 endA recA1 uidA(ΔMluI)::pir.* | Invitrogen |

**Table S2**. List of plasmids used in this study

| Plasmid | Description | Source |
|---|---|---|
| pRK600 | *Tra* and *Mob* genes. *Ori ColE1*. $Cm^R$ | Lab collection |
| pTn7-M | *ori R6K*. Mobilizable Tn7 element with *aacC1* (gentamicin resistance). $Gm^R$ $Km^R$ | (3) |
| pTns-1 | *ori R6K*, Transposase (TnSABC+D) operon. $Ap^R$ | (8) |
| pBG25 | *ori R6K*. Mobilizable Tn7 element with *aacC1* (gentamicin resistance) and msfGFP, under the control of 14F promoter and a bicistronic linker. $Gm^R$ $Km^R$ | (3) |
| pBG-control | *ori R6K*. Mobilizable Tn7 element with *aacC1* (gentamicin resistance). $Gm^R$ $Km^R$ | This study |
| pBG25-control14F | *ori R6K*. Mobilizable Tn7 element with *aacC1* (gentamicin resistance) and an empty 14F promoter. $Gm^R$ $Km^R$ | This study |
| pBG25-mCherry | *ori R6K*. pBG25 carrying mCherry under the control of 14F promoter and a bicistronic linker. $Gm^R$ $Km^R$ | This study |
| pBG25-fpvAp-mCherry | *ori R6K*. pBG25-mCherry with *fpvA* promoter and a bicistronic linker. $Gm^R$ $Km^R$ | This study |
| pBG25- fpvBp-mCherry | *ori R6K*. pBG25-mCherry with *fpvB* promoter and a bicistronic linker. $Gm^R$ $Km^R$ | This study |
| pBG25-14FfpvB | *ori R6K*. pBG25 with *fpvB* gene. $Gm^R$ $Km^R$ | This study |
| MBP-1.0 | *Ori ColE2*. GFPmut3b under the control of constitutive Ptet. | (6) |
| MBP-swapped | *Ori ColE2*. mRFP1 under the control of constitutive Ptet. | (6) |
| pBG25-GFP/RPF-control14F | *ori R6K*. pBG-control14F with Ptet regulated GFP/RFP expression. $Gm^R$ $Km^R$ | This study |
| pBG25-GFP/RPF-control | *ori R6K*. pBG-control with Ptet regulated GFP/RFP expression. $Gm^R$ $Km^R$ | This study |
| pBG25-GFP/RFP-14FfpvB | *ori R6K*. pBG25-14FfpvB with Ptet regulated GFP/RFP expression. $Gm^R$ $Km^R$ | This study |

**Table S3**. Oligonucleotides used in this study

| Name | Use / Restriction sites | Sequence 5´-3´* |
|---|---|---|
| mChFWeco<br>mChREVbam | Cloning mCherry into pBG25 // EcoR1/BamH1 | FW:tgg**gaattc**ttAGCAAGGGCGAGGAGGATA<br>REV:aga**ggatcc**TTACTTGTACAGCTCGTCCATGCCGC |
| fpvApFWpac<br>fpvApREVavr | Cloning *fpvA* promoter into pBG25-mCherry // PacI/AvrII | FW:gtc**ttaattaa**AACTTTTTCATCCGGTTTTCCGTGGGTAAAACGTCATACTAAGAGAATTGAGAAGCGATCTCATTTATTCGCAACTTGCCAACCT**cctagg**gcc<br>REV:ggc**cctagg**AGGTTGGCAAGTTGCGAATAAATGAGATCGCTTCTCAATTCTCTTA |

| | | |
|---|---|---|
| | | GTATGACGTTTTACCCACGGAAAACCG GATGAAAAAGTTtt**aattaa**gac |
| fpvBpFWpac<br>fpvBpREVavr | Cloning *fpvB* promoter into pBG25-mCherry // PacI/AvrII | FW:act**ttaattaa**GGCTGCGCTCCGTGGGCG GGAT<br>REV: aat**cctagg**GCGGCGCAGGCAGGCCGTT |
| mChFWconf<br>mChREVconf | Cloning confirmation primers for pBG25-mCherry, pBG25-fpvAp-mCherry and pBG25-fpvBp-mCherry | FW:cgacctagggcccaagttcac<br>REV:ATGTTGACGTTGTAGGCGCCGG |
| fpvB+RBSFWxba<br>fpvB+RBSREVhind | Cloning *fpvB* with its natural ribosome into *pBG25* // AvrII-Xba/HindIII | FW:cgc**tctaga**GCCATCCAGGACACTGCAG ATG<br>REV:gcg**aagctt**TCAGAGCGAGTACTTCAC CGTG |
| fpvB+RBSFWconf<br>fpvB+RBSREVconf | Cloning confirmation primers for pBG25-14FfpvB | FW:ttacgaaccgaacaggctt<br>REV:cgccagggttttcccagtcacgac |
| GFP/RFPFW_MBPpac<br>GFP/RFPREV_MBPxho | Cloning of Ptet-GFP/RFP from MBP-1.0/ MBP-swap into pBG25-control14F // PacI/XhoI | FW:cac**ttaattaa**tccctatcagtgatagagattgacatccc<br>REV:cta**ctcgag**aacgcgaagtaatcttttcggttttaaag |
| GFP/RFPFW_MBPpac<br>GFP/RFPREV_MBPbam | Cloning of Ptet-GFP/RFP from MBP-1.0/ MBP-swap into pBG25-14FfpvB // Pac/BamHI | FW:cac**ttaattaa**tccctatcagtgatagagattgacatccc<br>REV:cgt**ggatcc**aacgcgaagtaatcttttcggttttaaag |
| BG25_C14FFWpac<br>BG25_C14FREVxho | pBG25-control14F backbone amplification to host Ptet-GFP/RFP from MBP-1.0/ MBP-swap // PacI/XhoI | FW:cac**ttaattaa**gcccgttgacatgacatggtt<br>REV:ctat**ctcgag**gacgtcttgacataagcctgttcgg |
| BG25_14FBFWpac<br>BG25_C14FBREVbam | pBG25-14FfpvB backbone amplification to host Ptet-GFP/RFP from MBP-1.0/ MBP-swap // Pac/BamHI | FW:cac**ttaattaa**gcccgttgacatgacatggtt<br>REV:tat**ggatcc**gacgtcttgacataagcctgttcgg |
| BG25_TAGFW<br>BG25_TAG_GFPREV<br>BG25_TAG_RFPREV | Cloning confirmation primers for plasmids carrying Ptet-GFP/RFP from MBP-1.0/ MBP-swap | FW:tttacgaaccgaacaggctt<br>REV(GFP):ACGGGAACTACAAGACACGT G<br>REV(RFP):CTGCGTGGTACCAACTTCCC |
| Tn7RFW<br>glmsDownREV | Confirmation of Tn7 element integration | FW:cacagcataactggactgatttc<br>REV gcacatcggcgacgtgctctc |

*Bold regions indicate restriction sites for enzymatic digestion. Lower case indicates intergenic regions.

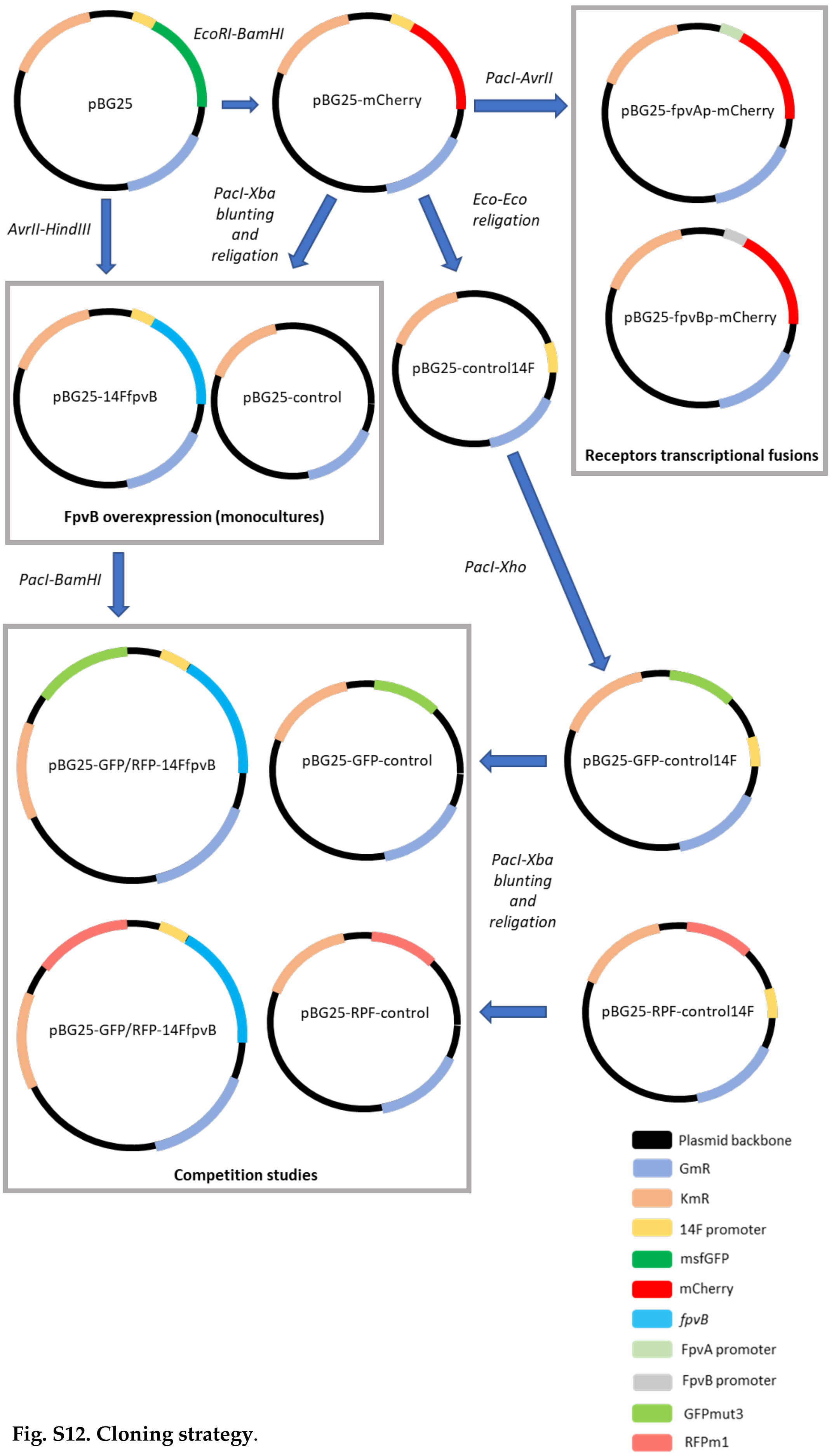


**Fig. S12. Cloning strategy**.

***Flow cytometry workflow***

All experiments were conducted in an Attune NxT flow cytometer equipped with acoustic focusing using the parameters summarised in Table S4. A summary of the gating strategy is presented in Fig. S13. Mixtures of GFP and RFP labelled strains were run as controls of the pipeline (Fig. S14). A representative example of a competition experiment is shown in Fig. S15)

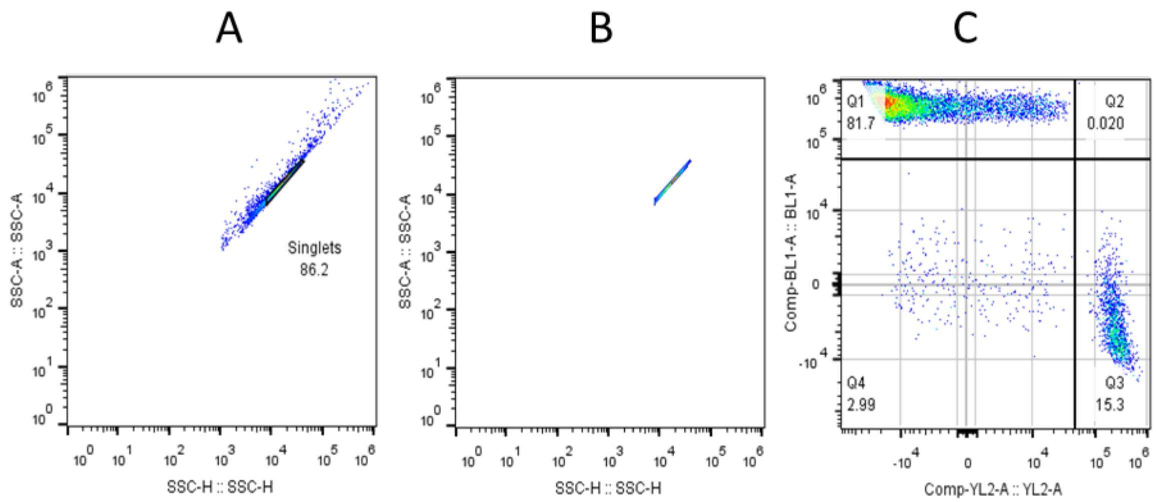


**Fig. S13. Example of gating strategy in a competition assay**. (A) The events corresponding to the population of interest (POI / singlets), were identified in a SSC-A vs SSC-H scatter plot and captured in a gate. This gate was preserved in all samples including the different biological replicates of the competition studies. (B) The captured POI represents (in this example) 86.2% of the total number of events. (C) The singlet gated POI in sectors. The majority of the cells belong to the GFP+ (Q1) and RFP+ (Q3) sub-populations.

**Table S4. Attune NxT parameter configuration**

| **Parameter** | **Voltage** | **Logic gate / Threshold** |
|---|---|---|
| FSC | 720 | AND / 10 x 1000 |
| SSC | 420 | AND / 1 x 1000 |
| BL1 Ex:488 nm Em:530/30 nm | 620 | - |
| YL2 Ex:561 nm Em:620/15 nm | 800 | - |

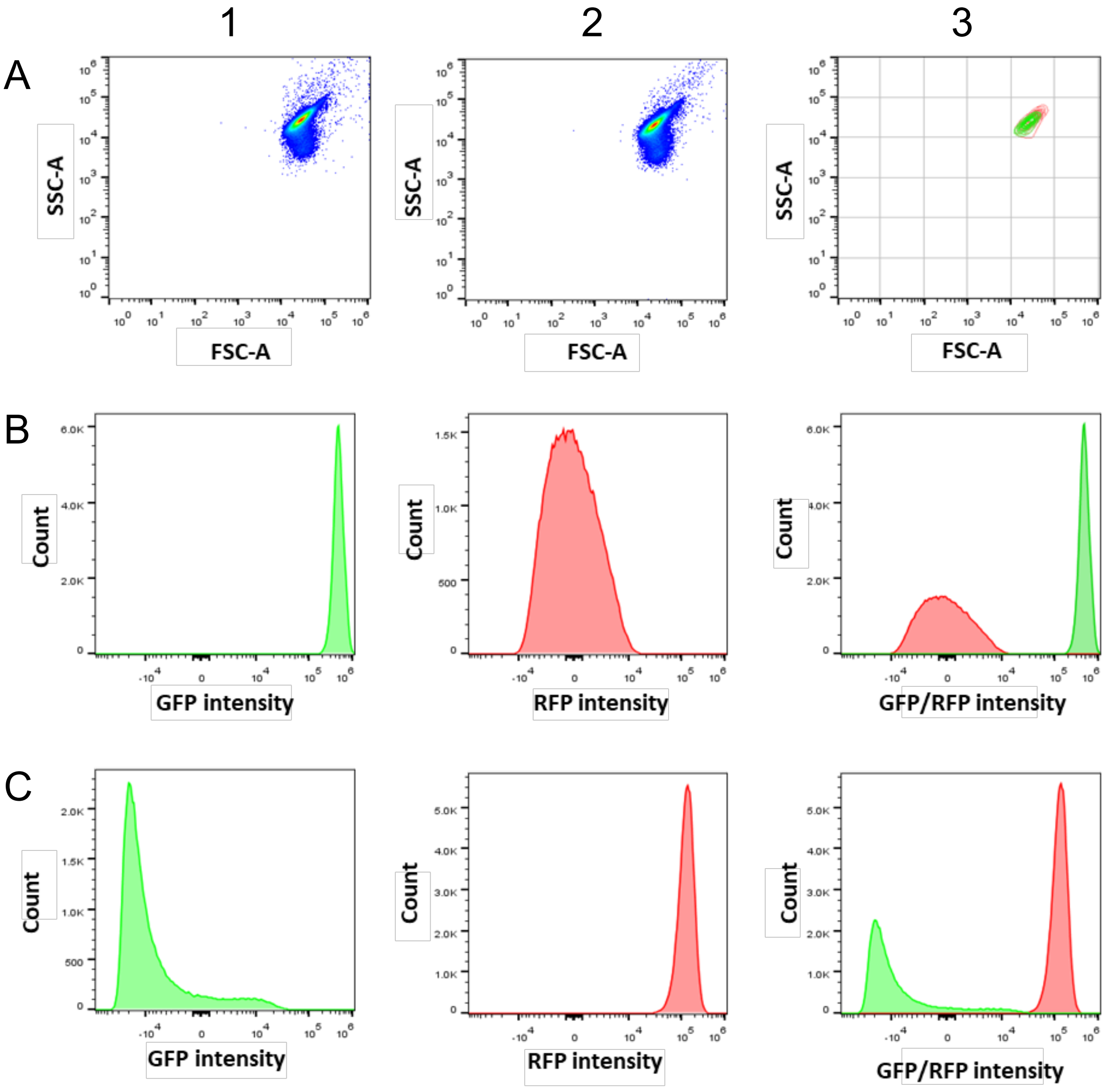


**Fig. S14. Analysis of a mixed population of GFP and RFP tagged MPAO1 bacteria**. A 0.5:0.5 mixture of cells was used as a control to test the FACS settings. Rows represent (A) SSC-A vs FSC-A scatter plots; (B) Histograms for GFP fluorescence (Ex:488 nm Em:530/30 nm); (C) Histograms for RFP fluorescence (Ex:561 nm Em:620/15 nm). Columns represent (1) GFP+ population; (2) RFP+ population; (3) GFP+ and RFP+ populations overlaid.

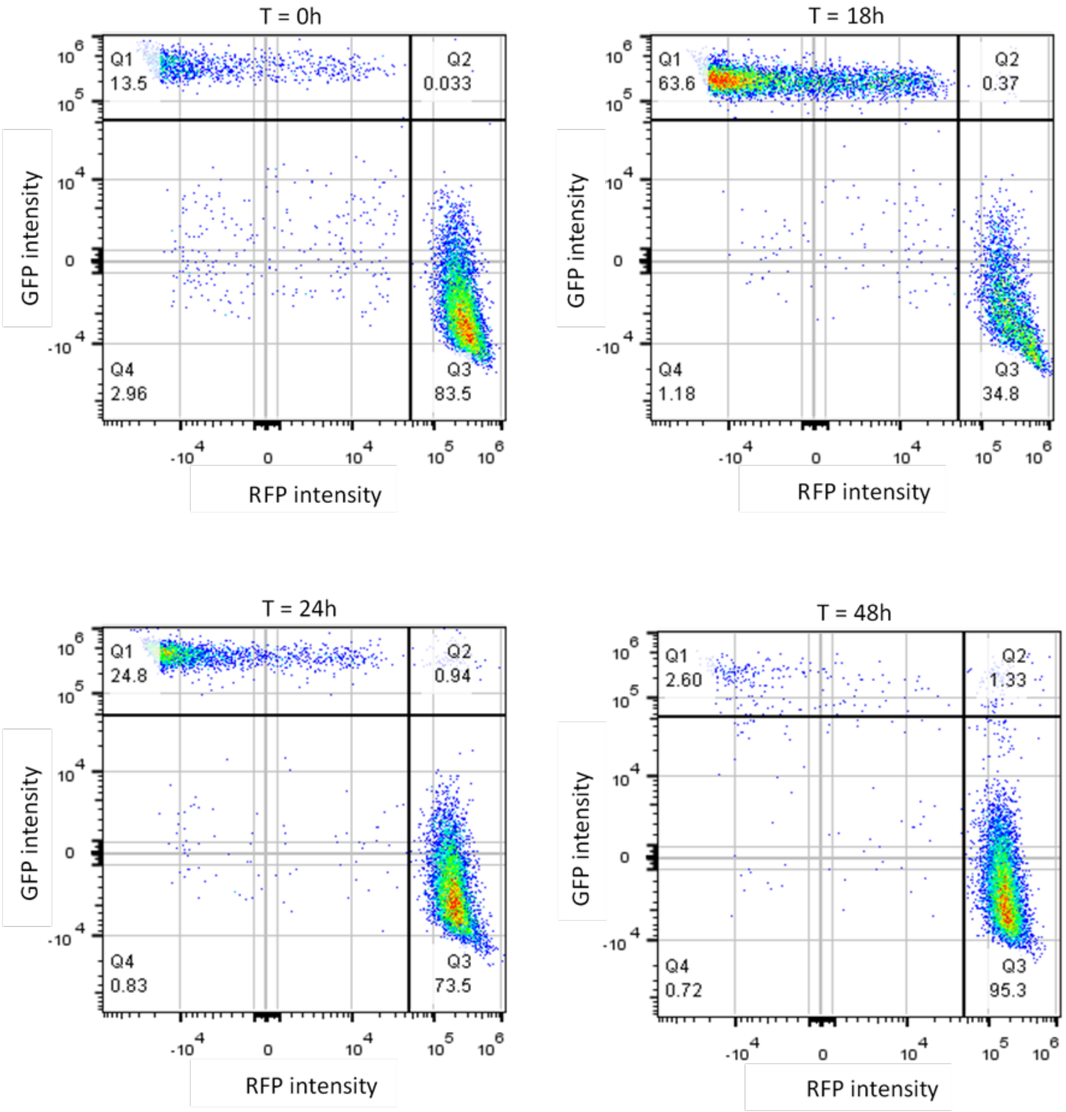


**Fig. S15. Population dynamics of a representative competition experiment monitored by FACS.** The example illustrates a competition assay between MPAO1 labelled with RFP (MPAO1::Tn7:*aacC1*:RFP; RFP+/GFP- Q3) and Δ*fpvA* (Δ*fpvA*::Tn7:a*acC1*:GFP; GFP+/RFP- Q1) in the presence of gentamicin. From top left to bottom right: 0, 18, 24 and 48 h. Values in the quadrants represent the frequency of each population within the singlets events. Horizontal axis represents red fluorescence corresponding to the RFP signal (MPAO1). Vertical axis represents green fluorescence corresponding to the GFP signal (Δ*fpvA*).
